# Radiation-Induced DNA Damage and Repair Effects on 3D Genome Organization

**DOI:** 10.1101/740704

**Authors:** Jacob T. Sanders, Trevor F. Freeman, Yang Xu, Rosela Golloshi, Mary A. Stallard, Rebeca San Martin, Adayabalam S. Balajee, Rachel Patton McCord

**Affiliations:** Department of Biochemistry & Cellular and Molecular Biology, University of Tennessee, Knoxville, TN 37996; UT-ORNL Graduate School of Genome Science and Technology, University of Tennessee, Knoxville, TN 37996; Radiation Emergency Assistance Center and Training Site, Cytogenetics Biodosimetry Laboratory, Oak Ridge Institute for Science and Education, Oak Ridge Associated Universities, Oak Ridge, TN 37830

**Keywords:** Genome Organization, DNA Repair, DNA Damage, Hi-C, Ataxia-telangiectasia mutated (ATM), Topologically Associating Domains (TADs)

## Abstract

The three-dimensional structure of chromosomes plays an important role in gene expression regulation and also influences the repair of radiation-induced DNA damage. Genomic aberrations that disrupt chromosome spatial domains can lead to diseases including cancer, but how the 3D genome structure responds to DNA damage is poorly understood. Here, we investigate the impact of DNA damage response and repair on 3D genome folding using Hi-C experiments on wild type cells and ataxia telangiectasia mutated (ATM) patient cells. Fibroblasts, lymphoblasts, and ATM-deficient fibroblasts were irradiated with 5 Gy X-rays and Hi-C was performed after 30 minutes, 24 hours, or 5 days after irradiation. 3D genome changes after irradiation were cell type-specific, with lymphoblastoid cells generally showing more contact changes than irradiated fibroblasts. However, all tested repair-proficient cell types exhibited an increased segregation of topologically associating domains (TADs). This TAD boundary strengthening after irradiation was not observed in ATM deficient fibroblasts and may indicate the presence of a mechanism to protect 3D genome structure integrity during DNA damage repair.

## INTRODUCTION

Ionizing radiation (IR) is a well-known carcinogen, and IR exposure inflicts a wide spectrum of DNA lesions among which the double strand break (DSB) is the most lethal lesion^1^. Mis-rejoining of DSBs leads to chromosome translocations, some of which may be oncogenic in nature. Therefore, efficient DNA repair is necessary to maintain genome integrity by suppressing chromosome translocations and genomic alterations^2^. Most studies thus far have focused on IR-induced DNA damage and repair at the level of the linear DNA sequence with a special emphasis on mutations, cancer induction, and downstream effects for cell death and organ/tissue damage. But, the human genome is not only a linear DNA sequence, but is also organized in a three-dimensional (3D) structure which influences the proper regulation of transcription, replication, and repair ^3, 4, 5, 6^. Considering the importance of 3D genome structure in various cellular activities, it is important to understand whether IR-induced DNA damage alters the genome architecture and whether the 3D genome architecture is restored after DNA repair.

Translocations, deletions, and other genomic aberrations that may follow DNA damage can lead to cancer by directly mutating genes or altering their regulation^7, 8^. Recently, it has become clear that the disruption of 3D genome domains can also be oncogenic^9^. Does the process of DNA repair protect the 3D folding of the genome as well as the linear DNA sequence? Certain cell types are considered to be more radiosensitive than others, but little is known about what contributes to their radiosensitivity. The possibility remains that cell type specific chromosome positioning, along with initial epigenetic chromatin and folding states can influence which translocations occur and how well DNA is able to repair after exposure to IR, helping to explain why certain cell types are more sensitive to radiation^8, 10, 11^.

Previous studies suggest that DNA repair efficiency may differ for heterochromatin and euchromatin ^12, 13^. Heterochromatic regions may be more mobile and move to DNA repair sites, where they decondense^14, 15^. Condensation or decondensation of specific chromatin regions may not be determined by their preexisting histone modifications^14^, suggesting that other factors may contribute to changes in 3D genome structure after DNA damage. One previous study demonstrated spatial clustering of DSBs in active genes by inducing specific breaks and measuring their interactions with Capture Hi-C^16^. This suggests that changes in the structure of local genome domains may happen at a broader scale after IR. Additionally, CCCTC-binding factor (CTCF) and cohesin have been shown to be early responders to DNA damage induced by IR^17, 18, 19, 20^. These proteins have also been recently demonstrated to play significant roles in chromosome folding^21, 22, 23, 24^, contributing to the formation of topologically associating domains (TADs). These genomic domains interact more frequently within themselves than with other regions and appear to help bring genes into contact with appropriate regulatory elements^25, 26^. The disruption of TAD boundaries can lead to developmental diseases and cancer^27, 28^. The prevailing model of TAD formation proposes that CTCF helps to establish TAD boundaries by blocking the loop extruding activity of cohesin^23^. Rapid recruitment of CTCF and cohesin to the sites of DNA damage may indicate that preservation of 3D genome architecture after DNA repair completion is important for genome integrity.

The above-mentioned studies suggest that changes occur in the 3D genome structure at multiple levels of its hierarchical organization in response to IR. Therefore, we sought to understand how IR and the subsequent repair of DNA influences genome organization across length scales in a cell type-specific manner. We employ chromosome conformation capture followed by high throughput sequencing (Hi-C) to characterize genome organization before and after exposure to X-rays (5 Gy)^29^. In a previous study, we used multi-color fluorescence in situ hybridization (mFISH) and low-resolution Hi-C experiments on human blood cells and lymphoblasts to examine changes in the contacts and positioning of whole chromosomes at 24 hours post-irradiation. We found that while individual cells harbored a variety of chromosomal translocations as observed by mFISH, at a population level, no recurrent translocations were evident in the Hi-C data^7^.

Here, with additional cell types and repair mutants at more timepoints and higher resolution, we assay the cell type-specific effects of IR on the hierarchical layers of genome organization. Hi-C was performed on fibroblasts and lymphoblastoids at 30 min and 24 h after exposure to 5 Gy X-rays. Changes in the overall genome organization were observed in the irradiated human fibroblasts and lymphoblastoid cells when compared to their non-irradiated counterparts. Human lymphoblastoid cells (GM12878) exhibited more evident changes in genome wide 3D organization when compared to fibroblasts. However, both cell types exhibited a strengthening of TAD insulation, suggesting that TADs were more segregated after X-ray exposure. This effect persisted at 5 days post-IR. To determine whether increased TAD insulation after IR is linked to the DNA damage/repair pathway, we examined the 3D genome organization changes in cells with compromised double strand break repair: human fibroblasts deficient in the ataxia telangiectasia mutated (ATM) gene. We observed that the TAD strengthening that occurs in several repair-proficient cell types after IR does not occur in ATM deficient cells, suggesting that this genome structure change is dependent on the ATM DNA repair pathway.

## RESULTS

### BJ-5ta skin fibroblasts exhibit subtle changes in genome architecture after X-ray irradiation

To determine how X-ray irradiation affects genome organization, we irradiated confluent BJ-5ta foreskin fibroblasts with 5 Gy X-rays and analyzed cells at 30 minutes and 24 hours post-irradiation. The majority of cells exposed to 5 Gy X-rays show chromosomal damage and translocations^7^. As cells at different cell cycle phases (G1, S and G2/M) respond differently to IR exposure, we grew the fibroblasts to a confluent, contact-inhibited state in which most cells are in G0/G1 phase (Supplementary Figure 1a)^30, 31^. To confirm the induction of DSBs by IR, immunofluorescence and western blot analyses were performed for phosphorylated-H2AX (γ-H2AX), a surrogate marker of DSBs^32, 33^. We observed that X-ray irradiated BJ-5ta cells showed an elevated level of γ-H2AX protein at 30 minutes post-exposure followed by a decline at 24 hours post-IR, indicating an efficient repair of DNA DSBs (Fig. 1a and Supplementary Figure 2a-b).

**Figure 1.**
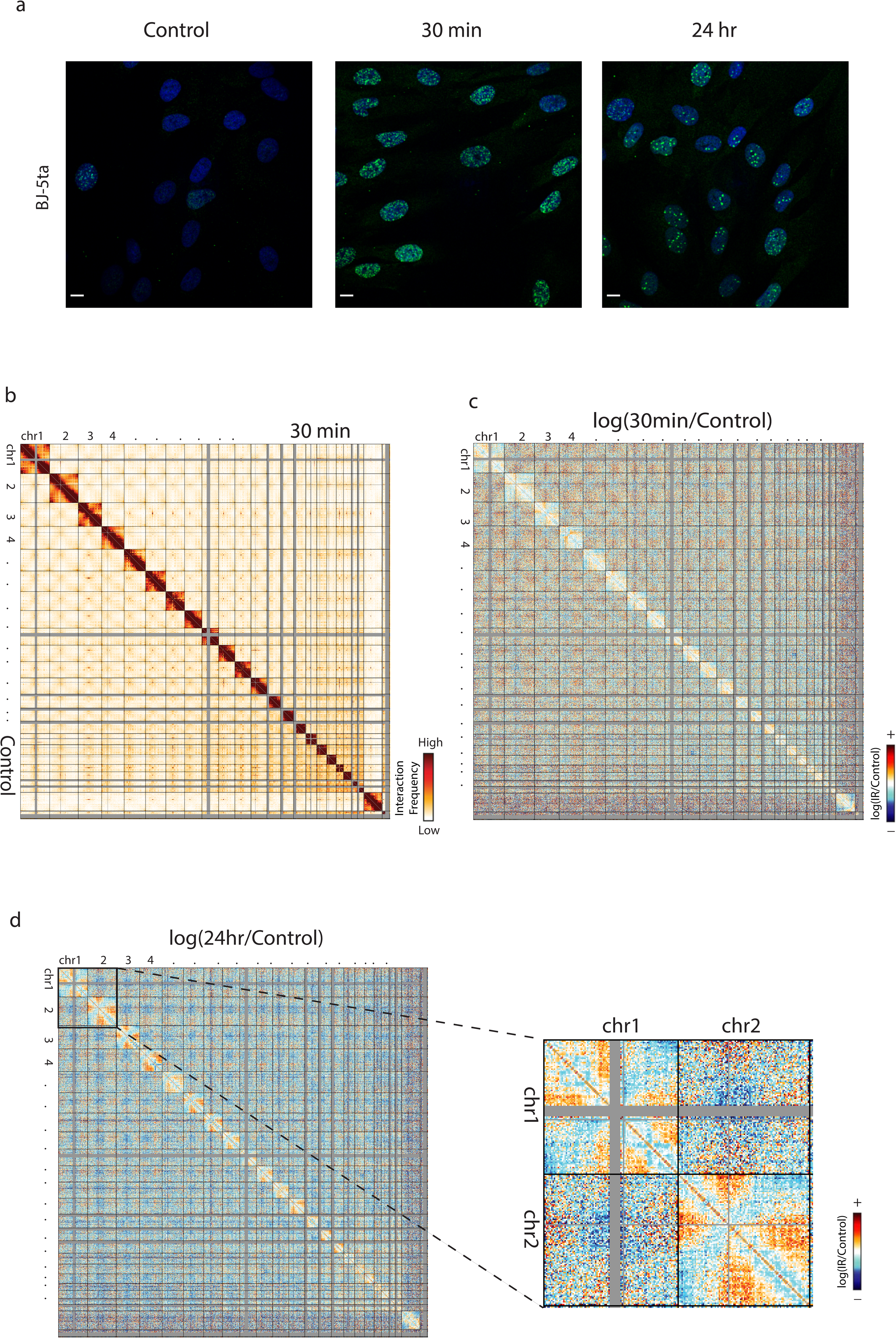
Subtle changes to genome organization in BJ-5ta skin fibroblasts exposed to IR. **a** BJ-5ta stained with yH2AX (green) and DAPI (blue) after before or after exposure to 5 Gy X-rays. Scale bar: 10um. **b** Genome wide contact heatmap for BJ-5ta non-irradiated (lower left) or 30 minutes after IR (upper right) in 2.5 Mb bins. **c** log(30 minutes post IR/Control) contact heatmap in 2.5 Mb bins. **d** log(24 hours post IR/Control) contact heatmap in 2.5 Mb bins. Inset reveals loss of interactions between telomeres on chromosomes 1 and 2.

To measure changes in genome-wide chromosome contacts after IR, Hi-C was performed immediately before IR exposure and at 30 minutes and 24 hours after exposure (see Methods and Supplementary Table 1 for statistics, processing, and replicate information). Strikingly, despite the observation of obvious DSB induction and repair events detected by γ-H2AX, only subtle changes were detected in whole genome contact maps after 30 minutes (Fig. 1b) or 24 hours (Supplementary Figure 3a) after IR exposure when compared with non-irradiated control cells. Chromosome territories were maintained at 30 minutes, and comparisons between 30 minutes and control show no obvious recurrent translocations, insertions, or deletions in the cell population. However, the log ratio of contact frequency 30 minutes post-irradiation compared to control shows a decrease in interactions between chromosome arms which suggests a separation of chromosome arms after irradiation (Fig. 1c). The decrease in interactions between chromosome arms persists at 24 hours, along with an even greater loss of interactions between telomeres (Fig. 1d). These experiments were also carried out in the BJ-1 hTERT cell line as an independent biological replicate. This is also an immortalized BJ fibroblast cell line made from a different clonal population than BJ-5ta. The BJ1 results cluster well with BJ-5ta experimental data (Supplementary Figure 4), and we observe similar changes to interaction patterns in these cells, specifically with the loss of interactions between telomeres within chromosomes (Supplementary Figure 3c), as well as changes at 30 minutes becoming more apparent at 24 hours post-IR (Supplementary Figure 3b-c).

### GM12878 lymphoblastoids exhibit evident genome-wide changes to chromatin architecture after exposure to IR

We next examined whether cell type specific differences exist in IR induced organizational changes by performing a comparative analysis on human fibroblasts and lymphoblastoid cells (GM12878). As stated before, X-rays induced DSB induction and repair were monitored at 30 min and 24 hours post IR exposure by measuring γ-H2AX using immunofluorescent and western blot techniques. As in BJ-5ta, an elevated level of γ-H2AX protein was observed in GM12878 cells at 30 minutes post-IR with a subsequent decline to baseline level at 24 hours post-IR (Figure 2a and Supplementary Figure 5a-b). We then performed Hi-C on GM12878 cells exposed to 5 Gy X-ray after 30 minutes and 24 hours. While Hi-C replicates showed high correlation (Supplementary Figure 4), we chose to treat replicates independently throughout subsequent analyses to provide independent corroboration of potential genome structure changes. Indeed, as discussed below, some features of contact changes were reproducible and others were not, suggesting that validating observations separately in multiple replicates is important, and that Hi-C datasets should not always be pooled immediately after a single reproducibility statistic is calculated.

**Figure 2.**
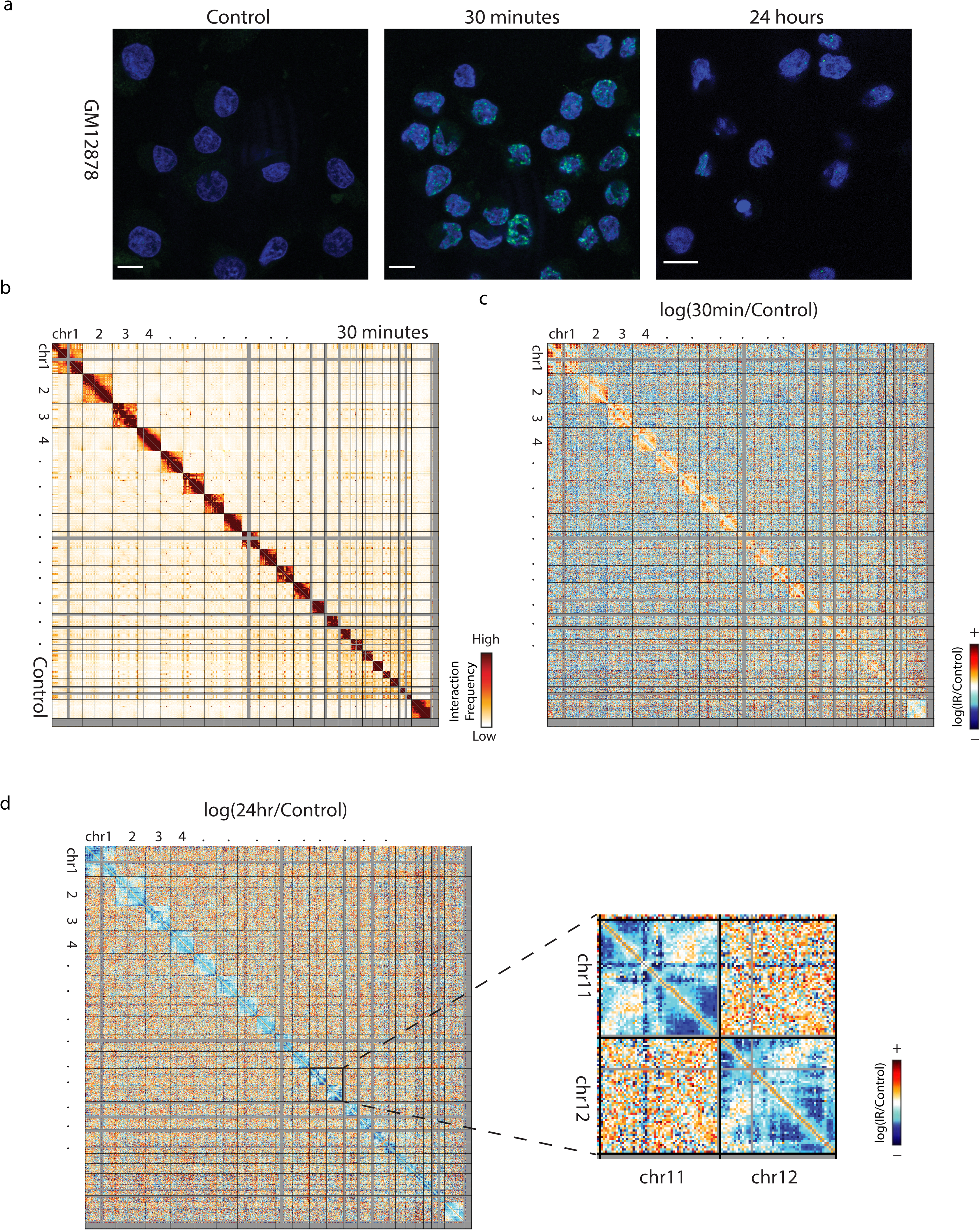
GM12878 exhibits evident changes to genome organization after exposure to IR. **a** GM12878 stained with yH2AX (green) and DAPI (blue) after before or after exposure to 5 Gy X-rays. Scale bar: 10um. **b** Genome wide contact heatmap of GM12878 in non-irradiated (left) or 30 minutes after IR (right) in 2.5 Mb bins. **c** log(30 minutes post IR/Control) contact heatmap in 2.5 Mb bins. **d** log(24 hours post IR/Control) contact heatmap in 2.5 Mb bins. Inset reveals example patterns of interactions that are lost across the genome after exposure to IR.

Much like the BJ-5ta cells, GM12878 did not exhibit any recurrent translocations, insertions, or deletions in genome structure (Fig. 2b and Supplementary Figure 6a). However, GM12878 cells exhibit a more variable response at 30 minutes post-irradiation. While there was an increase in the intrachromosomal interactions in both replicates, patterns of interactions observed were not consistent (Fig. 2c and Supplementary Figure 6b). Both replicates exhibited a loss of intrachromosomal interactions at 24 hours of post-exposure, and this loss of interactions was more pronounced in lymphoblastoid cells relative to fibroblasts. Also, GM12878 cells compared to fibroblasts showed more patterns of localized loss of intrachromosomal interactions at specific regions after 24 hours of IR exposure (Fig. 2d). These patterns of lost contacts were consistent between independent replicates. Interestingly, many of the regions which show increased interactions at 30 minutes decreased in interactions after 24 hours. Some of the regions involved in contact loss are notably gene dense, but further work will be needed to investigate whether other properties, like combinations of histone modifications, correspond to these regions of contact change after IR.

### Compartment identity is unchanged despite loss of distal interactions after IR exposure

Since we observed changes in intrachromosomal interactions in whole genome contact maps, we next examined higher resolution contact maps of individual chromosomes. These maps showed a depletion of interactions as distance from the diagonal increases after X-ray irradiation. This loss of distal interactions was confirmed by the generation of scaling plots which showed a loss of mid-to-long range interactions after 24 hours in both the BJ-5ta and GM12878 cell lines (Fig. 3a-b). The individual chromosome contact maps also showed no apparent changes in the plaid patterns between IR and control conditions, which suggests A/B compartment identity is not affected by IR^29^. The A and B compartment pattern captures the spatial segregation of euchromatin and heterochromatin within the nucleus. It has been previously shown that heterochromatin and euchromatin are differentially affected by DNA damaging irradiation and have different DNA repair responses^12, 13^. Surprisingly, we observed few instances of compartment identity switches (<∼1%) in both cell types and conditions as shown in both contact maps and eigenvector compartment tracks. (Fig 3c-f and Supplementary figure 7a-b).

**Figure 3.**
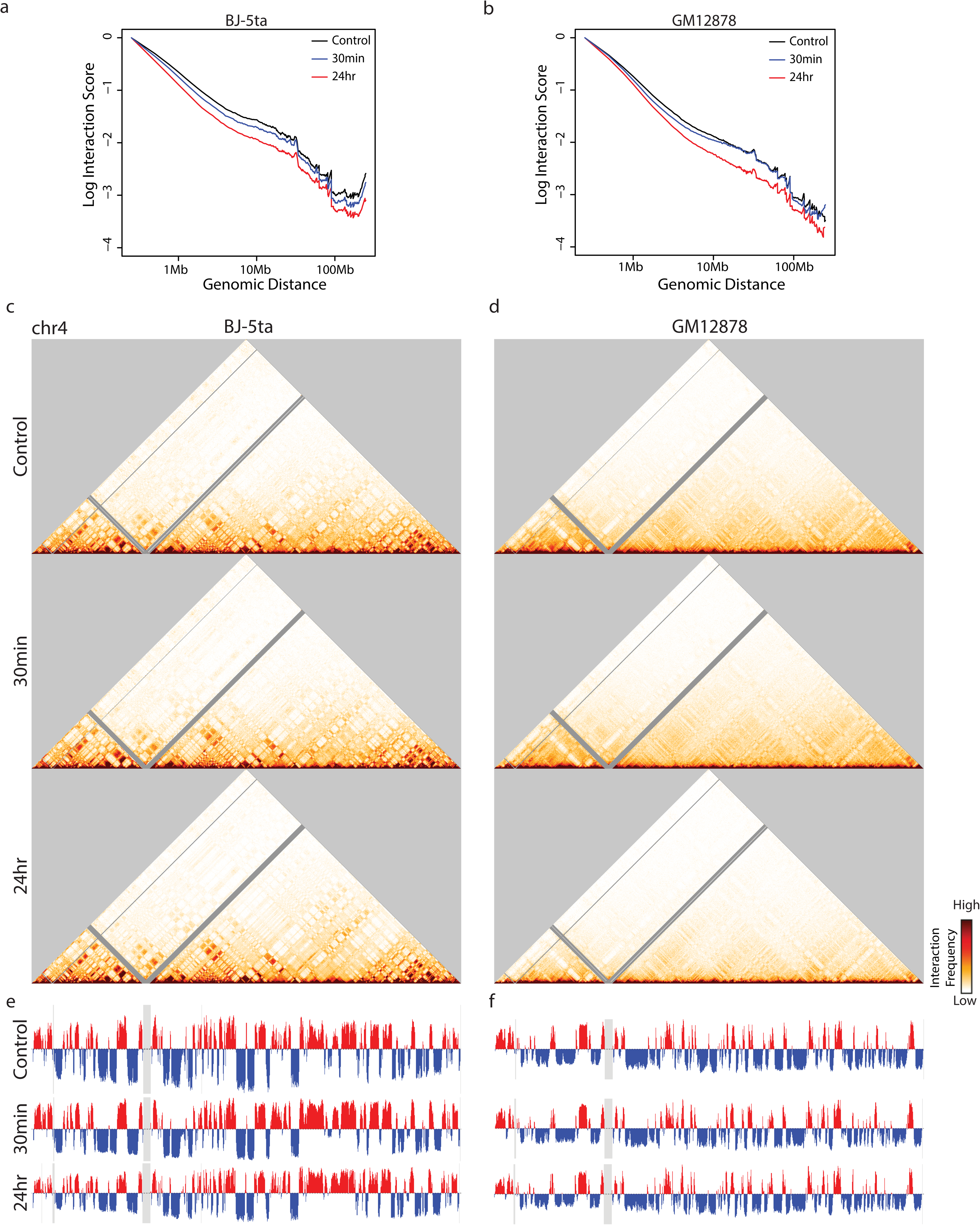
Healthy fibroblast and lymphoblast cell lines maintain compartments after exposure to IR. **a, b** Whole genome scaling plots at a 250 kb bin size in BJ-5ta (**a**) or GM12878 (**b)** cells for control, 30 minutes post IR, and 24 hours post IR. **c**, **d** 250kb binned Hi-C interaction heatmaps for chromosome 4 in BJ-5ta **(c)** and GM12878 **(d)** cells in control (Top), 30 minutes post IR (Middle), and 24 hours post IR (Bottom). **e,f** Plots of the first eigenvector from principal component analysis of chromosome 4 in BJ-5ta (**e**) and GM12878 **(f)** cells in control (Top), 30 minutes post IR (Middle), and 24 hours post IR (Bottom).

### TAD boundary strength and CTCF loops increase after exposure to IR

The chromosome architectural proteins CTCF and cohesin are recruited to sites of DNA damage^19^. CTCF occupancy increases both at binding sites flanking γ-H2AX domains^18^ and at sites of DNA lesions^17^ after DNA damage is induced. In addition to their role in the DNA damage response, CTCF and cohesin are known to be important for the formation of TADs, and alterations of CTCF and cohesin can lead to an overall change in local genome topology^34^. As previously mentioned, TADs have been identified as important for the spatial organization of gene regulation: enhancers and promoters are more likely to communicate within the same TAD and less likely to interact across TAD boundaries^27, 35^. The current model for TAD formation is that loop extrusion by the cohesin complex dynamically brings together genomic regions within TADs until extrusion is blocked by a boundary protein such as CTCF^22, 23^.

To assess whether TADs are affected after IR, we identified TAD boundaries and calculated the boundary strength (degree of spatial separation between TADs) for each boundary using both the InsulationScore^36^ and the Hicratio method^37^. Both methods determined that TAD boundary strength increases in both BJ-5ta and GM12878 cells 24 hours after IR (Fig. 4a-b and Supplementary Figure 8a-b).

**Figure 4.**
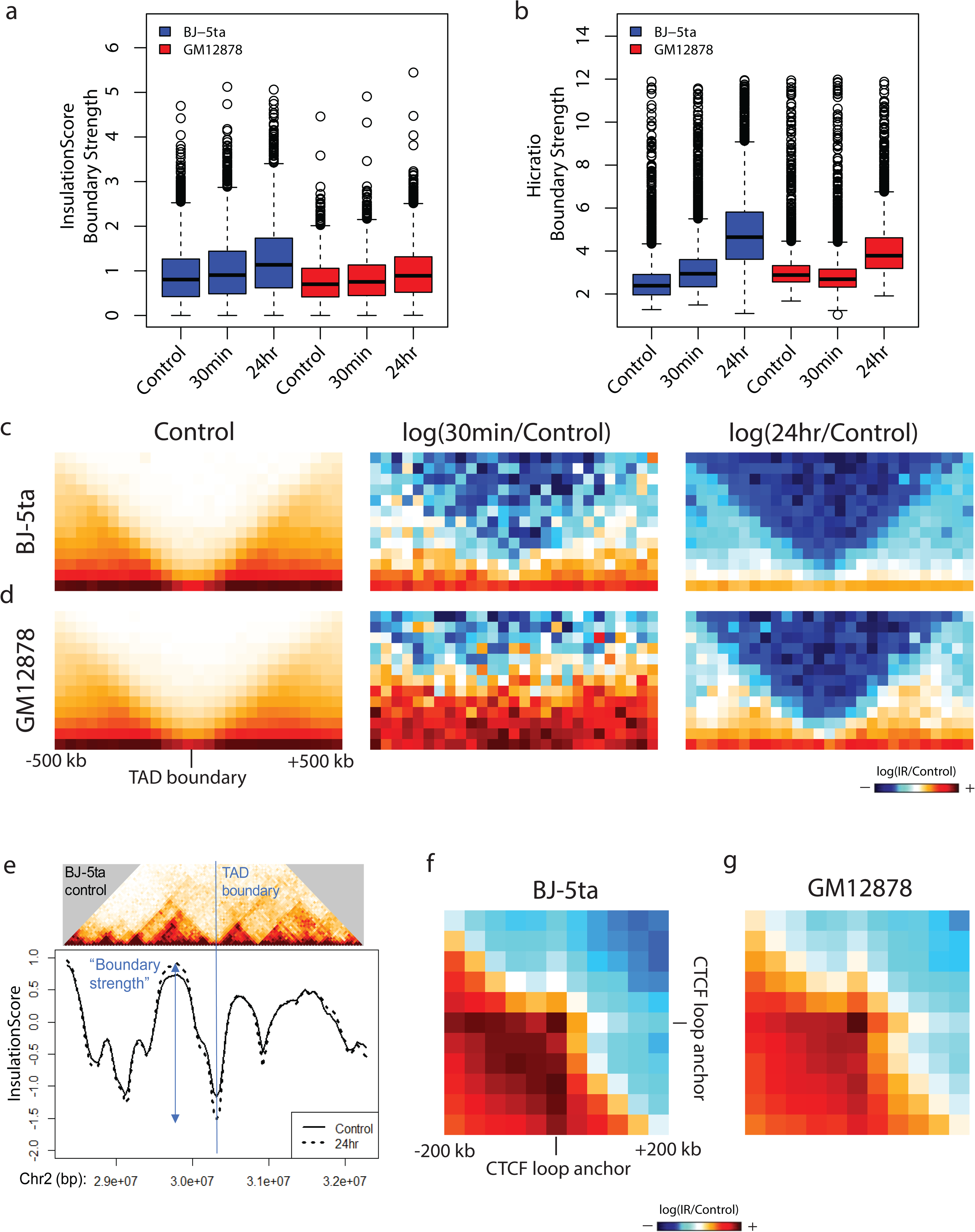
TAD boundary strength and CTCF loop strength both increase after IR. **a, b** TAD boundary strength boxplots calculated using both the InsulationScore (**a**) and Hicratio (**b**) methods for both BJ-5ta (blue) and GM12878 (red) cells. Boxes represent the upper and lower quartiles with the center line as the median. Upper whiskers extend 1.5×IQR beyond the upper quartile, and lower whiskers extend either 1.5×IQR below the lower quartile or to the end of the dataset. A Wilcoxon signed rank test with continuity correction determined that both BJ-5ta and GM12878 control boundary strength distributions were statistically less than their 30 minutes and 24 hour counterparts (p < 2.2X10^-16^). **c, d** Aggregate contact maps (40 kb bins) at TAD boundaries called by the InsulationScore method with strength greater than 1 in BJ-5ta (**c**) and GM12878 (**d**): Control (Left), log(30minutes/control) (Middle), and log(24hours/Control) (Right). **e** Example of strengthened TAD boundary. Top-BJ-5ta control Hi-C contacts around a TAD boundary from chr2. Bottom-InsulationScore profile across the region showing the sharper dip in 24 hr (dotted) vs. control (solid) at the TAD boundary. **f, g** Log ratio of 24 hours vs. Control aggregate interactions around loop anchors with CTCF sites (3,851 total loops) for BJ-5ta (**f**) and GM12878 (**g**).

One inconsistency that was observed between the two methods was the response at 30 minutes post IR in GM12878. In contrast to BJ-5ta, which had an increase in boundary strength in both methods after 30 minutes, GM12878 shows an increase as measured by the InsulationScore method but a decrease by the Hicratio method. This may be due to a subtle difference in the way that the algorithm detects TADs, so we focus on results that were consistent between both methods. To visualize the alterations in contacts around TAD boundaries, we plotted the average interaction profile around TAD boundaries called by the InsulationScore and Hicratio method (Fig. 4c-d and Supplementary Figure 9a-d). We observe a decrease in interactions across TAD boundaries after exposure to IR in all conditions except, again, in GM12878 at 30 minutes post IR. This divergence at 30 minutes between cell types may reflect different rates of DNA damage response and repair in different cell types, but the consistent effects on TAD strength at 24 hours across cell types and replicates suggests that this reflects a more fundamental feature of DNA damage response. The TAD boundary strengthening is observed not only in aggregate, but is also detectable at specific loci (Fig. 4e) Notably, we observe of a similar increase in TAD boundary strength 24 hours after IR in both lung fibroblasts (MRC-5) and an independent line of foreskin fibroblasts (BJ1-hTERT) (Supplementary Figure 8a-b). This supports the idea that the increase of TAD boundary strength is a general phenomenon, perhaps related to the dual roles of CTCF and cohesin in DNA damage response and genome architecture. The observation of this phenomenon in both fibroblasts and lymphoblasts also makes it less likely that the TAD strengthening observed is simply an effect of cell cycle state. BJ cells, as noted, are predominantly in G1/G0 phase before irradiation and become even a bit more uniform after irradiation. In contrast, GM12878 cells are still cycling at the time of irradiation and maintain similar distributions of cells in G1 and G2 at 24 h post-IR (Supplementary Figure 1b). Despite this difference, TAD strengthening is observed in both cell types.

If TAD boundaries are reinforced after IR because of increased recruitment of CTCF after DNA damage, we would expect that this might also increase contact frequency within CTCF-mediated loops. To test this hypothesis, we used the previously annotated set of loops from high resolution GM12878 Hi-C data, specifically analyzing those with CTCF motifs at both loop anchors^38^. We aggregated the interaction patterns across all loop regions and found that CTCF loops indeed show an increase in average strength after exposure to IR in both fibroblasts and lymphoblasts (Fig. 4f-g).

### ATM is necessary for IR-induced genome architecture changes

Changes observed in the 3D genome structure after IR exposure could be a passive side effect of DNA DSBs. However, the TAD boundary and CTCF loop strengthening that we observe are the opposite of what would be expected from a passive mixing of broken chromosomes, so we hypothesized that these changes are dependent on the cell’s active response to DNA damage. Ataxia telangiectasia mutated (ATM), a PI3K-like protein kinase, plays a key role in the repair of DSBs through phosphorylation of number of downstream targets ^39, 40, 41^. Patients with mutations in the ATM gene have an increased likelihood of several types of cancer, including breast cancer, leukemias, and lymphomas^42, 43^. ATM phosphorylates the histone variant H2AX which results in the recruitment of proteins involved in DNA damage response, including genome architecture proteins CTCF and cohesin^44^. Interestingly, the observation that γ-H2AX foci are bound by CTCF and are approximately the size of TADs has led several groups to hypothesize that ATM-mediated repair is restricted to TADs^45, 46^. To evaluate the role of ATM in the 3D genome response to X-ray irradiation, we performed similar IR experiments on ATM mutant fibroblasts immortalized with human telomerase reverse transcriptase (ATM-hTERT) as we did for previous cell types. Immunofluorescence imaging confirmed the faulty DNA repair pathway in these cells, shown by a lower mean fluorescence intensity of γ-H2AX (Fig. 5a and Supplementary Figure 10).

**Figure 5.**
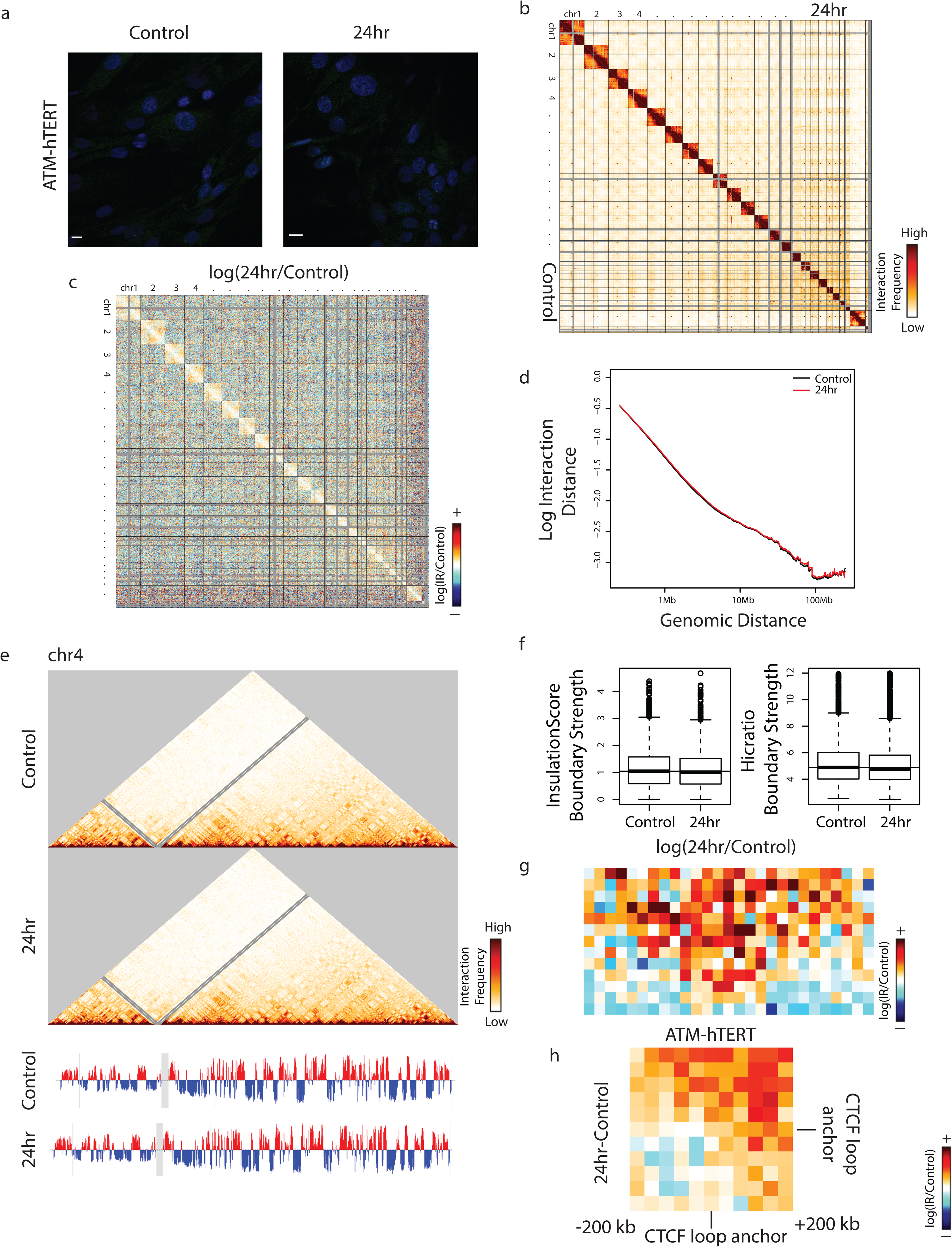
ATM is necessary to elicit 3D genome organization change after IR exposure. **a** ATM mutant cells stained with yH2AX (green) and DAPI (blue) after before after exposure to 5 Gy X-rays. Scale bar: 10um. **b** Genome wide contact heatmap of ATM mutant cells in non-irradiated (left) or 24 h after IR (right) in 2.5 Mb bins. **c** log(24 hours post IR/Control) contact heatmap in 2.5 Mb bins. **d** Whole genome log scaling plots at a 250 kb bin size in ATM mutant cells for control and 24 hours post IR. **e** 250kb bin size Hi-C interaction heatmaps for chromosome 4 in ATM mutant cells in control (Top) and 24 hours post IR (Bottom) show little difference in compartment as evidenced by little change to A (red) and B (blue) compartmentalization in the first principle component. **f** TAD insulation score boxplots calculated using both the InsulationScore and Hicratio methods for ATM mutant cells. A one-tailed Wilcoxon signed rank test showed that the 24 hour sample boundaries were not statistically increased compared to controls. **g** log(24hours/Control) heatmap of InsulationScore boundaries with strength greater than 1 averaged across the genome. **h** Subtraction of 24 hours and Control aggregate CTCF-anchored loop contact maps.

This contrasts with the results for BJ-5ta (Fig. 1a) and GM12878 cells (Fig. 2a), which both showed a higher mean intensity of γ-H2AX 30 minutes and 24 hours post irradiation. The presence of γ-H2AX at all after IR in the ATM mutants suggests that an alternate mechanism, likely the ATR pathway, in part compensates for the deficiency in ATM^47^.

We performed Hi-C on ATM-hTERT cells 24 hours post irradiation as a comparison to the condition in which the largest changes in genome organization occurred in BJ-5ta and GM12878 cells. Like both repair-proficient cell types, no recurrent translocations were observed in ATM-hTERT cells at 24 hours post IR (Fig. 5b). Additionally, the log ratio of 24 hours post-IR vs. control showed modest changes in contacts between conditions in these ATM mutant cells (Fig. 5c). We observed no change between IR condition and control scaling plots suggesting that the reduction of distal interactions in BJ-5ta and GM12878 cells is dependent on DNA damage response and repair. As in BJ-5ta cells, A/B compartment identities in ATM mutant cells are maintained 24 hours after exposure to IR (Fig. 5e). To evaluate TAD strength changes in ATM mutant cells after X-ray, we again used both the InsulationScore and Hicratio methods. In contrast to the increased TAD boundary strength in DNA repair proficient cells after X-ray, both methods showed a slight decrease in TAD boundary strength in ATM mutant cells after IR (Fig. 5f). We confirmed that there was an increase in interactions on average across TAD boundaries after X-ray in the ATM mutant cells by plotting the average interaction profile around TAD boundaries (Fig. 5g). Additionally, in contrast with BJ-5ta and GM12878, aggregate interaction patterns showed no increase of CTCF loops in ATM mutant cells (Fig. 5h). These results suggest that ATM is necessary for the increased segregation of TADs after IR, and that in the absence of ATM, genomic regions have a higher chance of interacting across TAD boundaries. To ensure that these results are attributable to ATM and not an artifact of TERT immortalization or specific to one cell line, Hi-C was performed in primary human fibroblasts derived from an AT patient (GM02052). We confirmed that in the absence of functional ATM, insulation between TADs remained the same or decreased. This is contrary to the TAD boundary strength increase in all the ATM-proficient cell lines (Supplementary Figure 8a-b).

### 3D genome response to IR is delayed in ATM mutants and persists in wild-type fibroblasts

The persistence of changes observed in the TAD strength at 24 hours after IR exposure in DNA repair proficient cells is striking, given that most DNA damage is repaired/misrepaired by this time point. Over an even longer time course after IR, cells that have failed to repair properly will often die and be eliminated from the population, selecting for cells that can continue to proliferate after repair. To determine the long-term effects of IR and the subsequent repair of DSBs DNA on the 3D genome, we irradiated BJ-5ta and ATM-hTERT fibroblasts with 5 Gy X-rays and then split the cells into sub-confluent densities after 24 hours. Cells were allowed to recover and proliferate until confluent (∼4-5 days). After performing Hi-C on the recovery condition from both cell lines, we observed that BJ-5ta fibroblasts maintained the loss of interactions at telomeric regions in large chromosomes observed at 30 minutes and 24 hours post-IR but showed a decrease in interactions at the centromeric regions among most chromosomes. Interactions that were observed to have changed after 24 hours post-irradiation are even more apparent in recovery (Fig 6a). Like 24 hours post-irradiation, the genome wide contacts of ATM mutant cells remained largely unaffected after 5 days recovery. The notable exception was that one replicate showed the emergence of two new translocations after 5 days of recovery: between chromosomes 17 and 21 and 19 and 21 (Fig. 6b). This is consistent with the repair-deficient nature of ATM mutant cells. As at 24 hours post-IR, mid-to-long range interactions remained decreased in BJ-5ta recovery as compared to the control (Fig. 6c). In addition, ATM mutants also had a decrease in mid-to-long range interactions after recovery (Fig. 6c), which differs from the response observed after 24 hours (Fig. 5d). A/B compartment identity is maintained in both BJ-5ta and ATM mutant cells after recovery, which is consistent with data obtained from 30 minutes and 24 hours IR responses (Fig. 6d).

**Figure 6.**
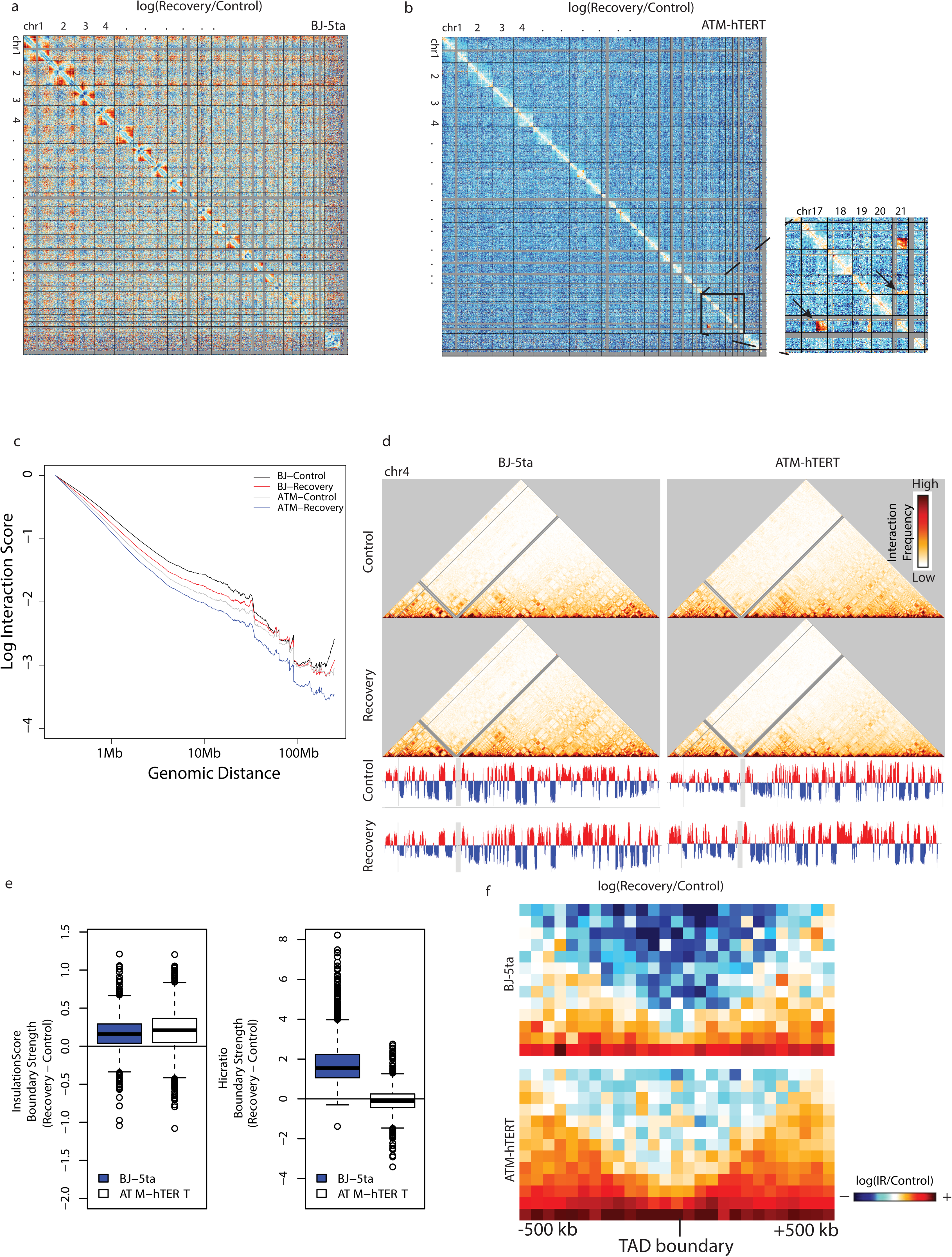
BJ-5ta maintains 3D organization changes after 5 days recovery, while ATM remains mostly unchanged. **a** log(Recovery/Control) contact heatmap in BJ-5ta binned at 2.5 Mb. **b** log(Recovery/Control) contact heatmap in ATM mutant cells binned at 2.5 Mb. Inset indicates translocations observed between chromosomes 17 and 19 and 19 and 21 at 5 days post IR that were not in the control ATM mutant cells. **c** Whole genome scaling plots comparing recovery and control samples at a 250 kb bin size in BJ-5ta and ATM mutant cells. **d** 250kb bin size Hi-C interaction heatmaps for chromosome 4 in BJ-5ta control (top left), BJ-5ta recovery (bottom left) ATM mutant control (top right) and ATM recovery (bottom right). PC1 compartment identity values remain relatively unchanged 5 days post IR. **e** TAD boundary strength boxplots calculated using both the InsulationScore and Hicratio methods subtracting recovery by control for BJ-5ta (blue) and ATM mutant cells (white). A Wilcoxon signed rank test with continuity correction determined that both BJ-5ta and ATM mutant control distributions were significantly lower than the recovery samples (p < 2.2X10^-16^) for InsulationScore. **f** log(Recovery/Control) heatmap of InsulationScore called boundaries with strength greater than 1 averaged across the genome in BJ-5ta (top) and ATM mutant cells (bottom).

TAD boundary strength measured by both InsulationScore and Hicratio methods remains higher than control 5 days after IR in BJ-5ta, suggesting that TADs maintain increased segregation long term after IR (Fig. 6e). Consistent with this result, the average TAD profile comparison showed a decrease in interactions across BJ-5ta TAD boundaries in 5-day recovery vs. control (Fig. 6f). Interestingly, ATM mutant cells also show some evidence of increased segregation of TADs after 5 days of recovery. The effect is not as strong or consistent, however, and varies by the quantification approach used (Fig. 6e). This again suggests that IR effects on TAD boundaries in ATM mutant cells are not the same as in repair-proficient cells but indicates that there could be a longer-term convergence, in which the ATM mutant cells that survive radiation long-term eventually show increased boundary strength. (Fig. 6f and Supplementary Figure 8d).

## DISCUSSION

The 3D architecture of the genome influences various DNA metabolic activities, including DNA repair to ensure the maintenance of genomic stability. Although IR induced gene mutations and chromosomal alterations have been well studied in various cell and animal model systems, IR-induced structural changes at the level of higher order chromatin organization have been given less attention.

In this study, we have identified notable and lasting effects of irradiation on genome organization within the cell population. Overall, the changes in 3D genome contacts that we observe after irradiation provide a new platform from which to consider the interplay between local chromatin structure, genome architecture proteins, TAD strength, and DNA repair after irradiation. Changes that are evident in population-averaged Hi-C data must have been occurring consistently, or at least in a substantial minority of cells, across the cell population. Our analysis methods also required that changes occur consistently across completely independent biological replicates. On the one hand, this is a potential weakness in the sense that we do not see the changes that are only occurring in certain individual cells, nor can we link a given change to a certain specific location of DNA damage. But, this requirement for signals to rise above the inherent variability in each cell means that the changes we observe report consistent features of the genome structure’s response to irradiation. Such features are more likely to reflect biologically controlled responses than would incidental effects of a certain site of damage in an individual cell.

We found that the effects of the same dose of radiation on genome organization vary between different cell types, causing different specific patterns of interaction change. This observation relates to key questions about the human health implications of radiation: What factors determine the downstream severity of radiation impacts (radiosensitivity) of exposed cells and tissues^48^? Why are certain cell types, genetic profiles, or disease states more radiosensitive than others^49^? Under highly similar culture and IR exposure conditions, we see that fibroblasts show an overall loss of chromosome arm contacts after IR while lymphoblasts show loss of interactions between specific compartment-scale genomic regions. We hypothesize that variations in initial chromosome structure and local chromatin state are a good candidate to explain some of the variations in the response to IR of different cells under different conditions. Since notable variations in initial chromosome structure occur across different cell types, in disease, and in response to different external conditions^50, 51^, our results support future work investigating the contribution of initial chromatin state to the changes we observe.

Most strikingly, this study revealed that an increase of TAD boundary strength after exposure to IR is consistent across cell types and that this phenomenon is dependent on functional ATM activity. Considering this new observation in the light of previous research suggests that this TAD strengthening after IR may arise from CTCF and cohesin’s dual roles in genome architecture and DNA damage response. Cohesin has long been known to be important in the DNA damage response and is a target of ATM^20, 52^. ATM signaling also increases CTCF recruitment to chromatin, CTCF sites are vulnerable to DNA breaks, and CTCF depletion increases genome instability^17, 53^. Meanwhile, increasing evidence supports the idea that cohesin and CTCF participate in the formation of TADs by loop extrusion^3, 23, 24, 34^. Thus, we propose that more cohesin complexes recruited to sites of DNA damage bring regions within TADs together more frequently as they extrude loops. As they do so, they would make H2AX phosphorylation within TADs more likely, potentially explaining the apparent coincidence of γH2AX foci and TADs^14, 45^. The extruding complexes would then be blocked by increased CTCF recruitment, further decreasing interactions between TADs and leading to the boundary strength increases we see across cell types.

Why would increased TAD strength matter biologically to the cells exposed to radiation? Such increased genome structure segregation in response to DNA damage could be an important part of the cell’s ability to maintain genome integrity and prevent deleterious translocations. It is known that translocations which occur across TAD boundaries can result in the misregulation of genes and be oncogenic, and it is known that translocation frequency is influenced by pre-existing genome contact frequency^8, 9^. Thus, decreasing contacts across TAD boundaries in response to DNA damage would be one way to decrease the frequency of translocation events that disrupt TAD boundaries. We observe long-term persistence of TAD strengthening after IR (even at 5 days after IR, when cells have been allowed to proliferate). Notably, the cells that survive to divide are those that have not been eliminated by mitotic catastrophe due to deleterious translocations. If TAD strengthening is protective, we would predict that exposure to and recovery from one round of IR would make the cell population more resistant to deleterious effects of a second round of IR. While it is difficult to quantitate what degree of boundary strength change would have a meaningful biological impact, previous work has suggested a role for similar changes in boundary elements in biological processes such as gene dosage compensation^36^.

Future work will be needed to clarify the role of chromatin state and genome architecture proteins in the responses to IR that we observe. Perturbations of chromatin state before IR exposure would reveal the impact of different initial structures on downstream effects of radiation. Similarly, measuring the effect of irradiation on cells with altered levels of CTCF or cohesin will determine whether these factors indeed affect the spread of γ-H2AX. Our data supports a model in which ATM is required for the reinforcement of 3D genome organization after exposure to IR. But, the possibility remains that not only ATM, but also the subsequent signaling cascade involved in DNA repair downstream of ATM may be necessary to elicit the genome structure responses we observe in healthy cells. To understand more clearly how DNA damage pathways influence the 3D genome architecture, the role of proteins such as DNA ligases and ATR must be interrogated more thoroughly^54, 55, 56, 57, 58, 59^.

Additionally, future work is needed to identify the biological importance of TAD strengthening after IR. It is possible that the TAD structure effects we observe are indeed a byproduct of repair processes, but not an important or controlled component of protecting the genome from aberrations. In the future, artificially altering TAD strength, by altering concentrations of architecture proteins, would allow a more direct perturbation of the effect of TAD strength on cellular radiosensitivity and downstream effects of radiation.

## METHODS

### Cell culture

Fibroblast cell lines include: BJ-5ta (ATCC, CRL-4001), BJ1-hTERT, MRC-5 (kind gift from Tim Sparer, University of Tennessee, Knoxville, Knoxville, TN), AG04405 (ATM-hTERT) (kind gift from Peter Lansdorp, University of British Columbia, Vancouver, Canada), and GM02052 (Coriell). The lymphoblastoid cell line GM12878 (Coriell) was also used. Information about cell lines and media formulations is summarized in Supplementary Table 2. All growth media and fetal bovine serum was purchased from Corning, except for GM12878 growth media. All fibroblast lines were passaged at a density of 80%. For irradiation experiments, fibroblast lines were grown to confluency prior to X-ray exposure. For the GM12878 suspension cell line, cells were passaged at a density around 500,000 cells per 1 mL of medium. For irradiation experiments, GM12878 cells were grown to a density of 1 million cells per 1 mL of medium prior to X-ray exposure.

### X-ray irradiation

Cells were irradiated with 5 Gy X-rays using a RadSource RS-2000 Biological Irradiator. X-rays were emitted with a concentration of 2Gy/min with cells being exposed for 2 minutes and 30 seconds (160.9 kV; 25.1 mA). After irradiation, cells were returned to an incubation chamber and kept at 37°C with 5% CO_2_. Cells were collected at 0 minutes, 30 minutes, or 24 hours post-irradiation and fixed for the respective downstream application.

### Immunofluorescence staining, image acquisition, and image analysis

Fibroblast cells were seeded on poly-D-lysine (PDL) coated 35 mm coverslip dishes prior to X-ray exposure. For GM12878 cells, coverslips were sterilized and coated in 0.1 mg/mL PDL. GM12878 cells were X-rayed while in suspension and centrifuged onto coverslips at 800×*g* for 10 minutes immediately before fixation. At the appropriate time points, all cells were fixed in 4% formaldehyde for 20 minutes at room temperature, washed in PBS for 5 minutes three times, and then concurrently permeabilized and blocked in blocking buffer (10% goat serum, 0.5% triton in 1X PBS) for 1 hour at room temperature. The primary antibody to γ-H2AX (Abcam-ab26350, 1:500) was diluted in a 1:1 mixture of blocking buffer and PBS, and incubated overnight at 4°C. Samples were then washed three times in PBS for 5 minutes and incubated in secondary antibody (Alexa Fluor 488 goat anti-mouse, Invitrogen; R37120) for 30 minutes at room temperature, in the dark. Samples were washed as previously described and exposed to 0.1 mg/mL DAPI for one minute, followed by washing. All PBS was removed by aspiration and samples were briefly air dried. Coverslips were mounted on slides with Slow fade Diamond Antifade Mountant (Molecular Probes, S36963).

Images were acquired using a Leica SP8 confocal microscope equipped with a 63× oil immersion objective (Leica) using the same settings. Image quantification was carried out by measuring the mean fluorescence intensity of γ-H2AX inside the nucleus divided by area of the nucleus in ImageJ. An ordinary one-way ANOVA test was used to determine statistical significance for all image quantification.

### Flow cytometry and cell cycle analysis

Cells were fixed before or after irradiation and fixed with 70% ethanol for 30 minutes at 4°C. Cells were then centrifuged for 10 minutes at 800×*g* for 10 minutes and resuspended with Guava® Cell Cycle Reagent (Luminex, 4500-0220). 10,000 events were collected for each condition on an Attune Nxt Acoustic Focusing Cytometer. Cell cycle analysis was done using Flowing Software 2.5.1 (Terho, P. Cell Imaging Core, Centre for Biotechnology, University of Turku, Finland).

### Hi-C

To prepare the Hi-C libraries and prepare for sequencing, at the appropriate time points, Hi-C experiments were performed according to the protocol detailed in Golloshi et al.^60^. Briefly, 5-20 million cells were fixed with 1% formaldehyde, suspended in cell lysis buffer for permeabilization, and homogenized by douncing. Crosslinked chromatin was digested overnight with HindIII. Sticky ends were filled in with biotin-dCTP, and the blunt ends of interacting fragments were ligated together. DNA was purified by two phenol-chloroform extractions and ethanol precipitation. Biotin-dCTP at unligated ends was removed, and the DNA was sheared to a target size of 200-400 bp by a Covaris sonicator (Covaris, M220). DNA between 100-400 bp was selected for using AMPure XP beads (Beckman Coulter). Biotinylated DNA was pulled down using streptavidin coated magnetic beads and prepared for multiplex sequencing on an Illumina platform using the NEBNext Ultra II kit (NEB). All end preparation, adaptor ligation, and PCR amplification steps were carried out on bead bound DNA libraries.

Sequencing was carried out at the Oklahoma Medical Research Foundation Clinical Genomics Facility on an Illumina HiSeq 3000 platform with 75 bp paired end reads or a NovaSeq 6000 with 50 bp paired end reads. Sequencing reads were mapped to the reference human genome hg19, filtered, and iteratively corrected using the pipeline detailed in Imakaev *et al.*^61^ and available on github (dekkerlab-cMapping). All processed Hi-C contact maps were scaled to a sum of 1 million to allow comparisons between conditions for subsequent analyses.

### Hi-C Reproducibility Assessment

A spearman correlation-based metric was used to assess reproducibility across all Hi-C datasets. First, the log of iteratively corrected interaction counts, binned at 2.5 Mb across the entire genome, was calculated and used to calculate correlation maps for each individual matrix. Each entry in such a Hi-C data correlation matrix represents the correlation between the full row and column of data at that position in the matrix. This is like the calculation typically performed to emphasize interaction patterns over the average distance decay for compartment analysis. Then, each genome-wide correlation matrix was converted to a vector, and all vectorized Hi-C matrices were compared with pairwise Spearman correlation to obtain the final reproducibility metric between each pair of datasets. The results from this approach match well with the published GenomeDISCO approach^62^.

### Interaction vs. Distance Scaling Plots

Using 250 kb binned iteratively corrected and scaled contact matrices, we extracted contact frequencies between bins at each genomic distance, excluding the diagonal bin (zero distance). We then used a loess fit to find a smooth curve describing interaction decay vs. distance. Loess curves for each chromosome were averaged together to generate a single scaling curve representing the whole genome. The interaction frequencies were then normalized to set the maximum value (loess fit value for the minimum distance) for each dataset to 1 and then plotted on a log scale.

### Compartment analysis

Principle components analysis was performed using the matrix2compartment perl script in the cworld-dekker pipeline available on github on 250 kb binned matrices. The eigen1 value, which typically represents the compartment profile, was used to determine A/B compartments.

### TAD calling, insulation contact maps, and TAD boundary score boxplots

Topologically associating domains (TADs) boundaries were called on 40 kb binned matrices using two methods: the InsulationScore method, as detailed in Crane et al.^36^, available as matrix2insulation in the cworld-dekker github package, and the Hicratio method, as described in Gong et al.^37^. For the InsulationScore, a 500 kb insulation square size was used, and for the Hicratio method, we used d = 400 kb.

Insulation contact maps in main figures were generated by averaging all called TAD boundaries with an insulation score greater than one as determined by the InsulationScore method. We plotted 500 kb away from the diagonal and 500 kb on either side of the TAD boundary. To remove biases introduced by the contact matrix diagonal, we excluded the 30 kb closest to the diagonal.

Boxplots were generated using boundary strength scores determined by both InsulationScore and Hicratio methods. All TAD boundaries were plotted for the InsulationScore boxplots. For Hicratio boxplots, outliers were removed by filtering out TAD boundaries with a Hicratio score greater than twelve or less than one.

### Western blot

At the appropriate time points, cells were trypsinized using standard protocols, and up to 5 million cells were collected by centrifugation (1000 rcf, 5 min, room temperature). After carefully removing the media from the pellet, cells were resuspended in 300 ml of RIPA buffer (Thermo Scientific, I89900), supplemented with protease inhibitors/EDTA (GenDepot 50-101-5485), and phosphatase inhibitors (GenDepot 50-101-5488). Cells were lysed by repeated pipetting followed by incubation, on ice, for 10 minutes. DNA was degraded by adding 1.5 μl of micrococcal nuclease (0.5 U/ml stock, Thermo Scientific, FEREN0181) and 3 μl of calcium chloride (100 mM stock) to each lysate, followed by incubation at 37°C for 15 min. Nuclease was inactivated by subsequent incubations at 68 °C for 15 min, and ice for 10 min. Cell debris was removed by centrifugation at 14,000 rcf for 15 minutes. Protein samples were stored at −80 °C until use. Protein concentration was measured using the BCA Protein Assay Kit. Thermo (234225), according to the manufacturer’s instructions.

Denatured protein samples were resolved using a mini gel tank (Invitrogen, A25977) for 35 min at 200V, using 4-12% Bis-Tris Plus gels (Invitrogen, NW04120BOX) for all other targets. Protein was transferred to low fluorescence PVDF membranes using the mini Bolt Blotting System (Invitrogen, B1000), along with system-specific reagents. Blotting was performed using the Odyssey TBS blocking system (Licor, 927-400000), according to the manufacturer’s protocol. Briefly, membranes were activated with methanol after transfer (1 min, RT), washed twice with milliQ water (5 min), twice with TBS (2 min), and blocked with Odyssey TBS blocking solution for 1 hour at RT. Primary antibodies were diluted in blocking buffer (containing 0.2% Tween), and incubated over night at 4°C as follows: γ-H2AX (1:1000, mouse monoclonal [9F3], Abcam). Beta actin was used as a loading control, using the appropriate antibody (1:10,000; rabbit polyclonal PA1-16889, Thermo Fisher or 1:10,000; mouse monoclonal MA1-140, Thermo Fisher) concomitantly with targets of interest. Detection was carried out using the following secondary antibodies: goat anti-rabbit (1:10,000; IRDye 680RD [92568071]; Licor) and goat anti-mouse (1:10,000, IRDye 800CW [95-32210]; Licor). Secondary antibodies were diluted in Odyssey TBS blocking buffer containing 0.2% Tween and 0.01% SDS. Membranes were incubated in secondary antibody for 1 hour at RT, followed by three washes (TBST, 5 min ea.).

Images of near infrared fluorescent signal were acquired using an Odyssey scanner (Licor) in both the red and green channels. Signal was quantitated using the Image Studio software (Licor).

## Data Availability

All relevant data supporting the key findings of this study are available within the article and its Supplementary Information files or from the corresponding author on reasonable request. The Hi-C data generated in this study have been deposited in Gene Expression Omnibus (GEO) under accession number XXXX.

## Code Availability

The majority of software and code used for the analyses presented are available as public github packages, as described in the Methods section. Any minor custom analysis scripts will be made available upon request.

## Acknowledgements

We thank Dianne Trent and Stephen Kania in the Biomedical and Diagnostic Sciences department of the University of Tennessee Veterinary Medical Center for advice and assistance with flow cytometry and Mariano Labrador for helpful discussions. This work was supported in part by a grant from Oak Ridge Associated Universities Directed Research and Development (ODRD) to A.B. and R.P.M.

## Author Contributions

R.P.M., A.B., and J.T.S. devised the project. Hi-C experiments were performed by J.T.S, T.F., and R.G. Image quantification and analysis was performed by M.S., R.S.M. and J.T.S. Western blot preparation and analysis was performed by R.SM. and J.T.S. Hi-C analysis was performed by J.T.S, T.F., Y.X., and R.P.M. The manuscript was prepared by J.T.S, T.F, and R.P.M, with input from A.B., R.SM, and R.G.

---

**Supplementary Figure 1.**
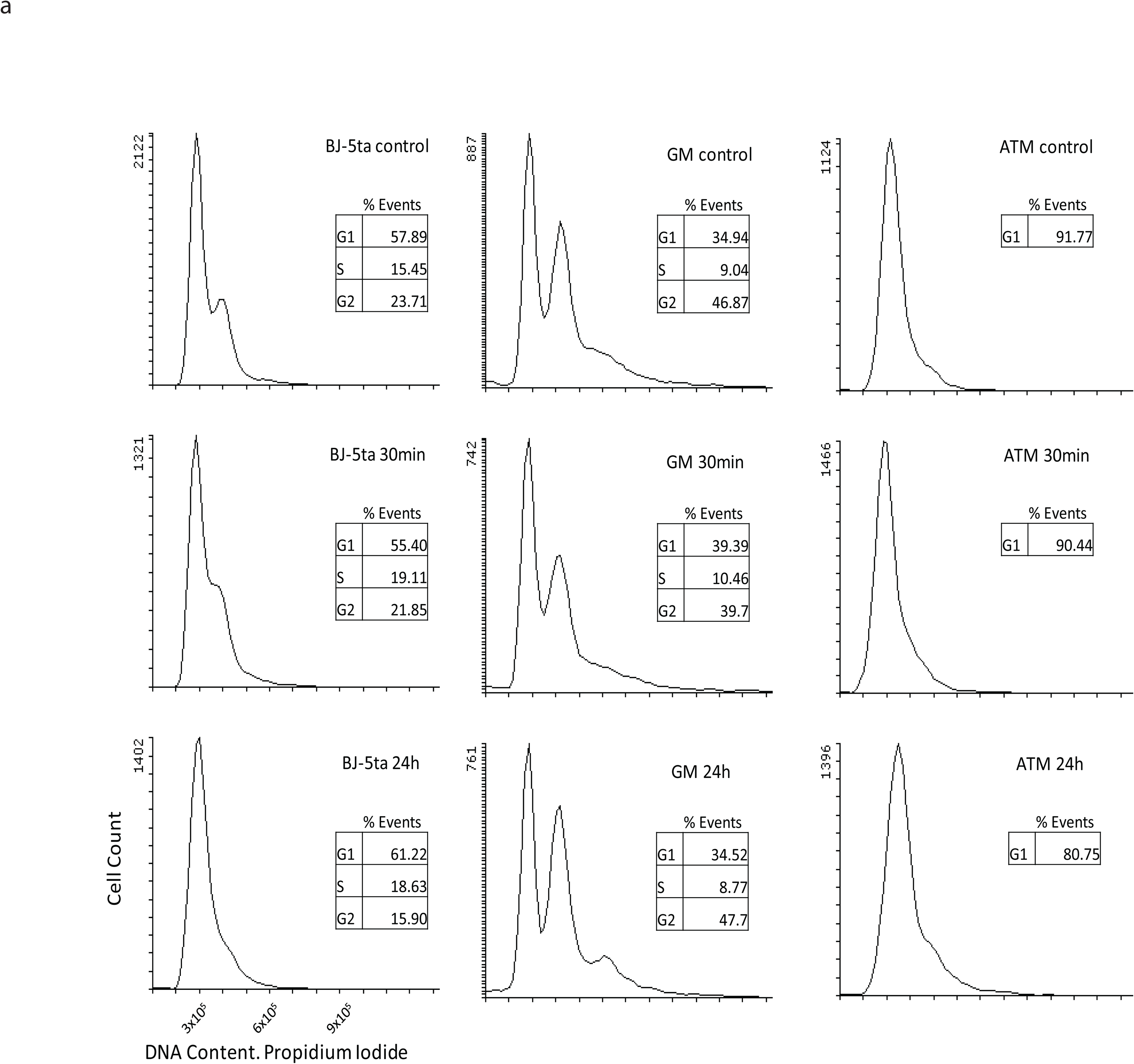
Flow cytometry analysis of BJ-5ta, GM12878, and ATM-hTERT cells before and after irradiation. Y-axis shows cell counts while x-axis represents the fluorescence measurement of DNA content. BJ-5ta cells are predominantly in G1 in all conditions but contain a small population of G2 in control and 30 minutes post IR. GM12878 cells maintain a very similar cell cycle state distribution before and after IR. ATM-hTERT cells are mostly G1 stalled in all conditions.

**Supplementary Figure 2.**
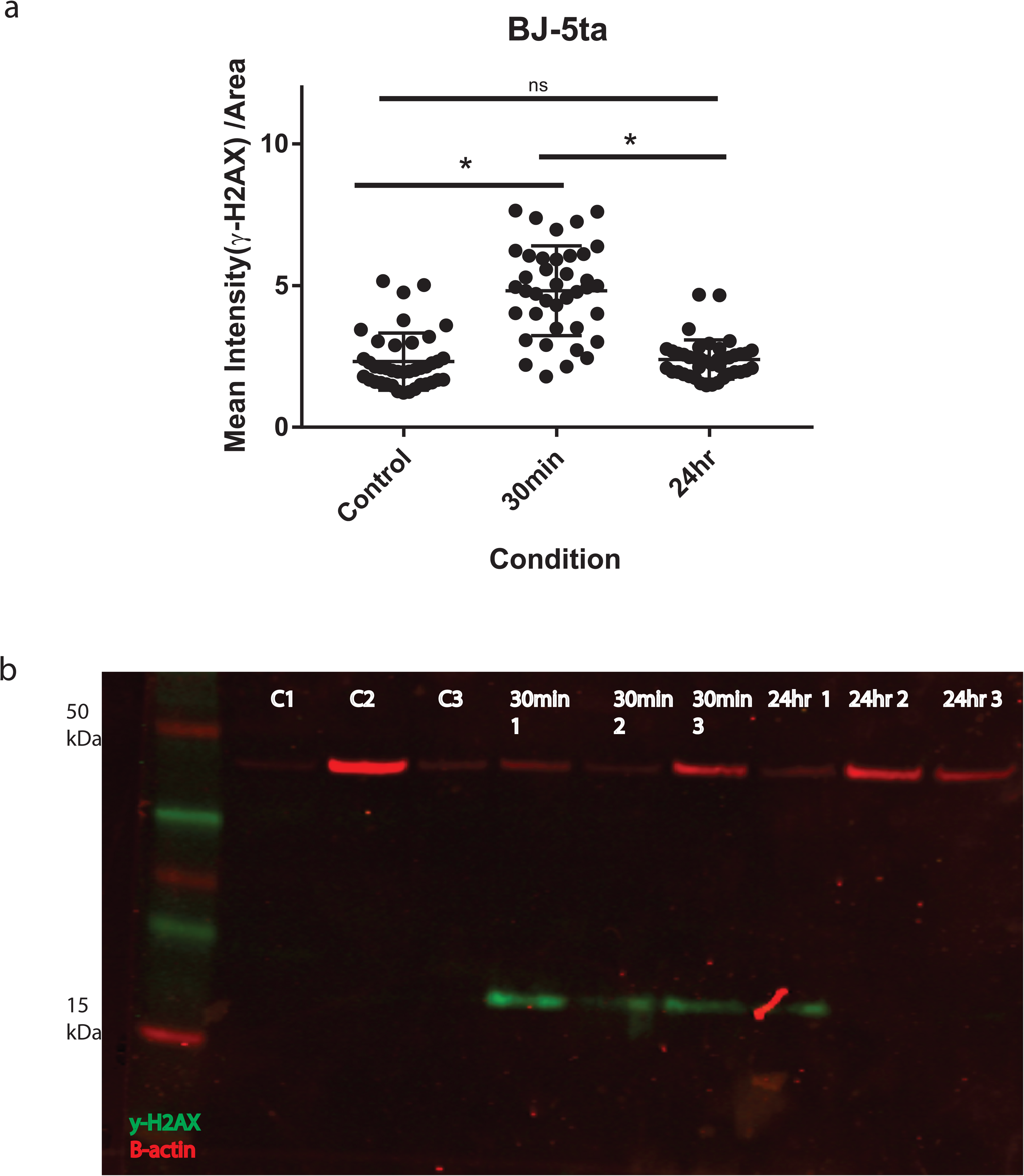
γ-H2AX levels increase in BJ-5ta after exposure to IR. **a** Quantification of mean fluorescence intensity in BJ-5ta cells after exposure to IR (n=40, * p <0.0001, one way ANOVA). **b** Western blot analyses for β-actin (red) and γ-H2AX (green). Bands around ∼15 kDa indicate the presence of γ-H2AX.

**Supplementary Figure 3.**
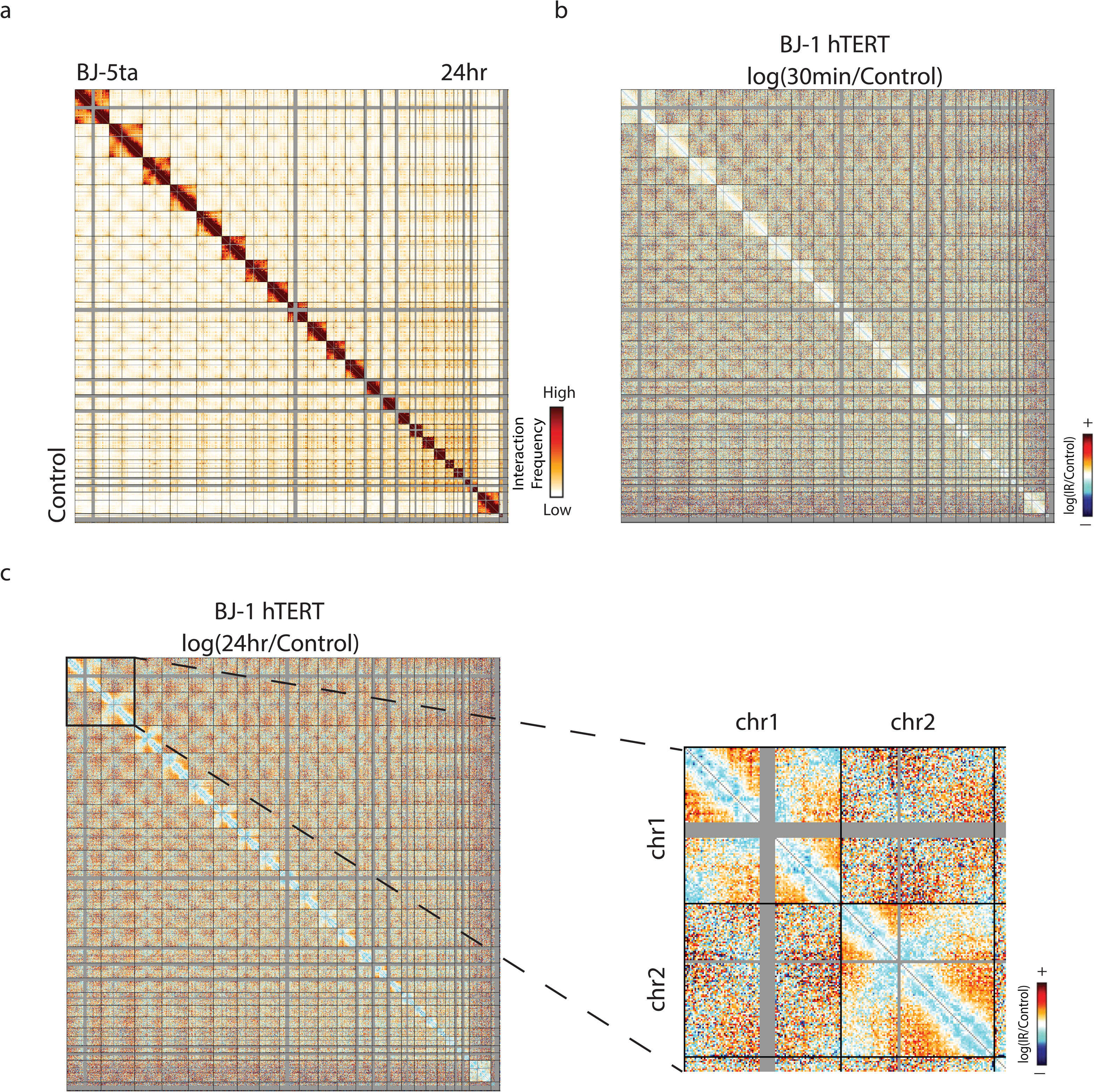
BJ-5ta and BJ1-hTERT share similar responses to IR. **a** Genome wide contact map of non-irradiated BJ-5ta (left) and 24 hours after irradiation (right). **b** log(30 minutes post IR/Control) BJ1-hTERT contact heatmap in 2.5 Mb bins. **c** log(24 hours post IR/Control) BJ1-hTERT contact heatmap in 2.5 Mb bins. Inset reveals loss of interactions between telomeres after IR in chromosomes 1 and 2.

**Supplementary Figure 4.**
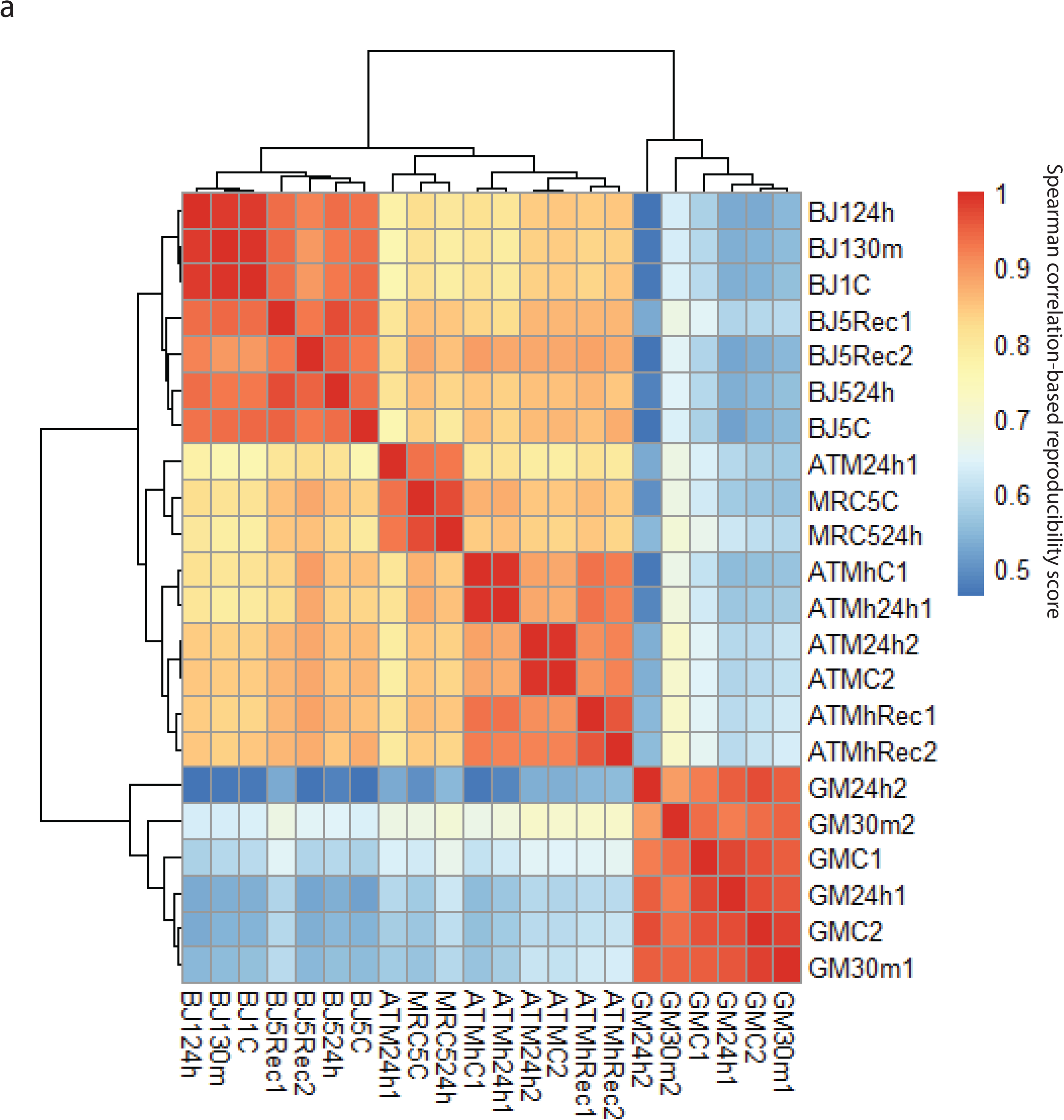
Reproducibility correlation plot for all Hi-C experiments. Spearman correlation-based reproducibility is calculated as described in the Methods. ATM proficient fibroblasts mostly cluster together and separately from ATM deficient fibroblasts. GM12878 cells cluster separately from fibroblasts. All replicates tend to cluster together rather than experimental conditions.

**Supplementary Figure 5.**
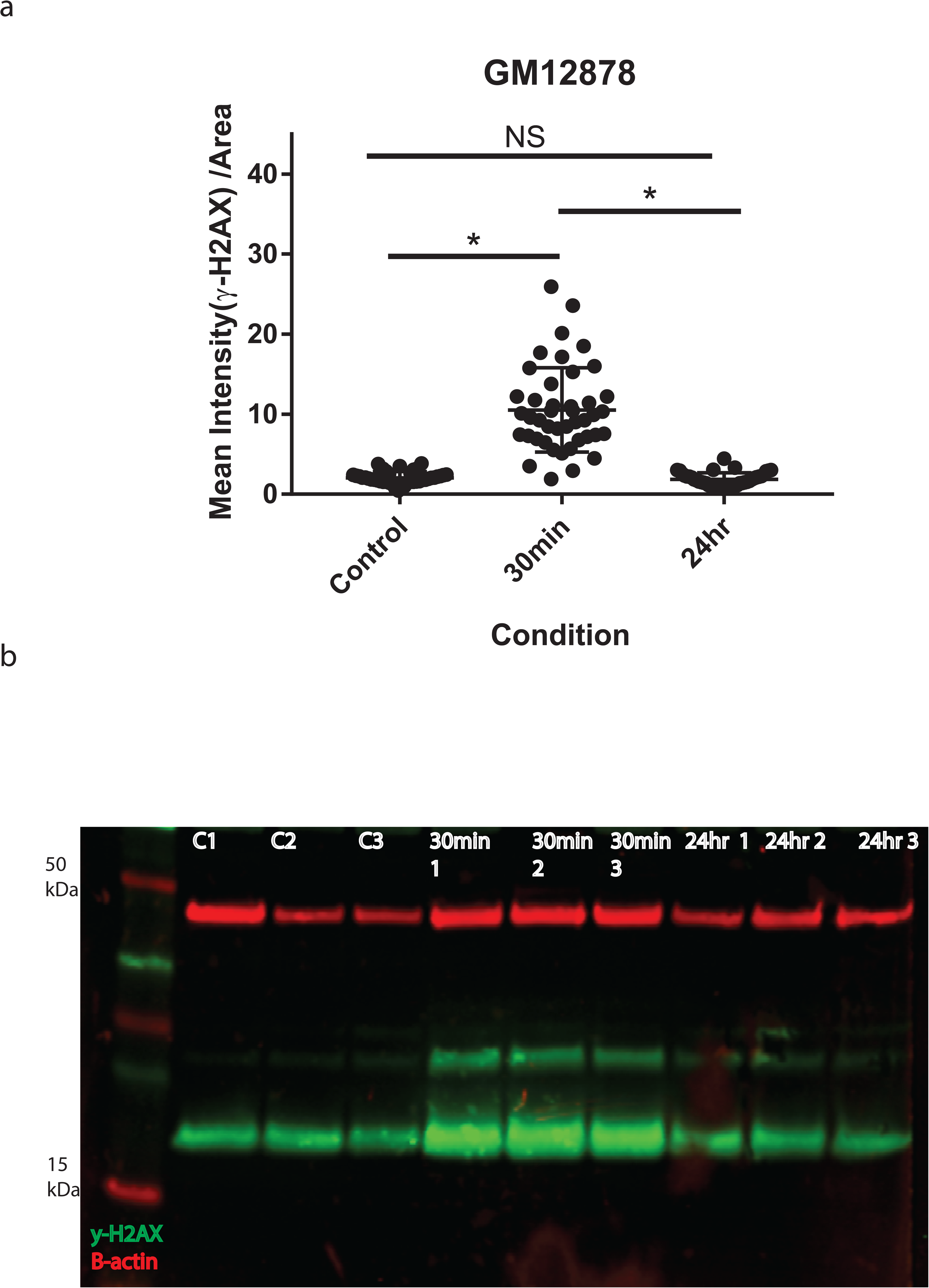
γ-H2AX levels increase in GM12878 after exposure to IR. **a** Quantification of mean fluorescence intensity in BJ-5ta cells after exposure to IR (n=40, p < 0.0001, one way ANOVA). **b** Western blot analyses for β-actin (red) and γ-H2AX (green). Bands around ∼15 kDa indicate the presence of γ-H2AX. GM12878 contains more γ-H2AX in control cells than BJ-5ta, but this may be due to cell division.

**Supplementary Figure 6.**
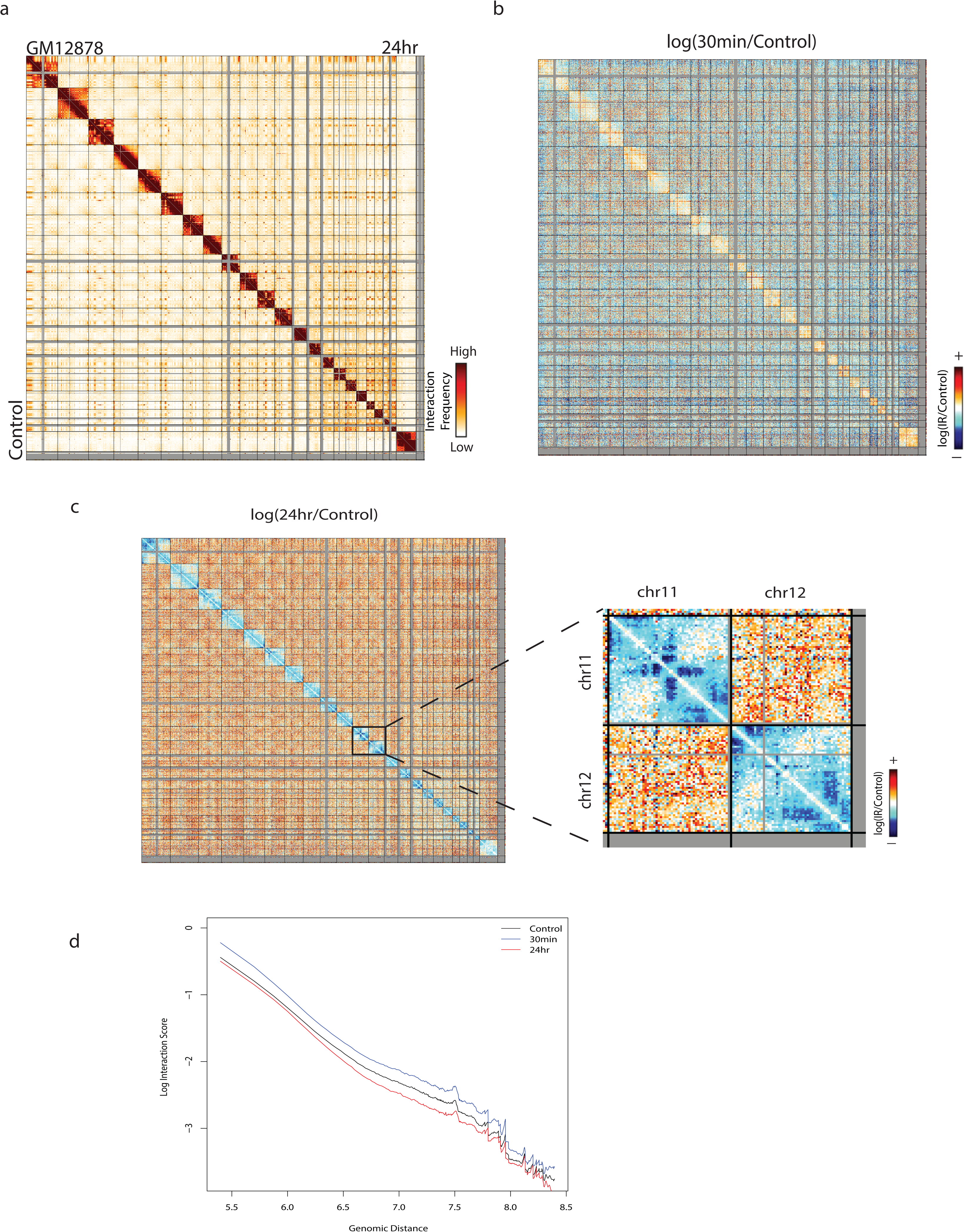
Similar major features of GM12878 genome structure change after IR are observed in a biological replicate (R1, R2 shown in main text figures). **a** Genome wide contact map of non-irradiated GM12878 (left) and 24 hours after irradiation (right). **b** log(30 minutes post IR/Control) GM12878 contact heatmap in 2.5 Mb bins. **c** log(24 hours post IR/Control) GM12878 contact heatmap in 2.5 Mb bins. Inset reveals similar specific changes within chr11 and chr12 as shown in main Figure 2. **d** Whole genome scaling plots at a 250 kb bin size in GM12878 (R1).

**Supplementary Figure 7.**
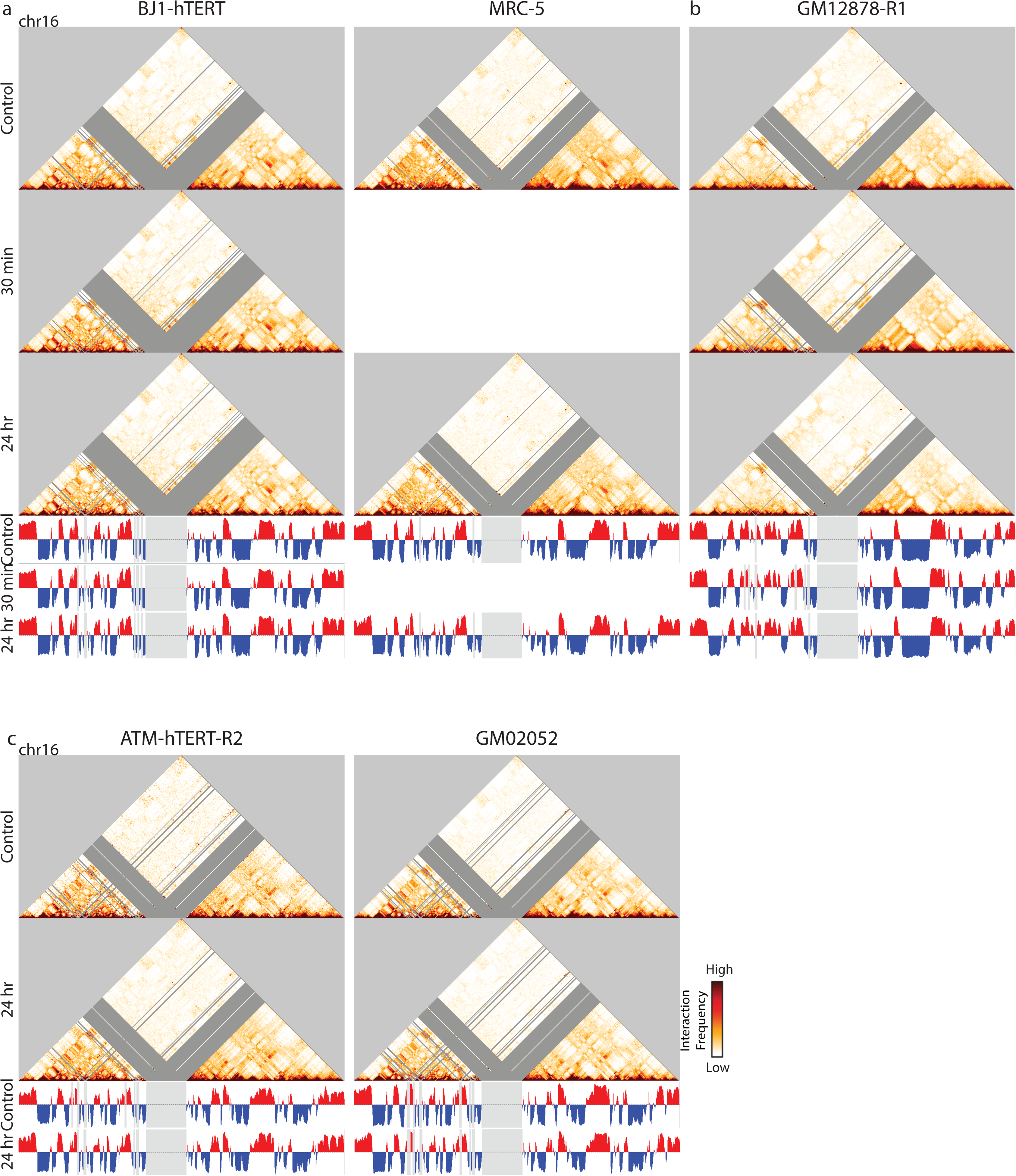
A/B compartment identity is robust to changes post-IR in healthy cell lines and ATM mutant cell lines. **a** 250kb bin size Hi-C interaction heatmaps (top) and plots of the first eigenvector from principal component analysis (bottom) for chromosome 16 in healthy fibroblasts: BJ1-hTERT (left) and MRC-5 (right). **b** 250kb bin size Hi-C interaction heatmaps (top) and plots of the first eigenvector from principal component analysis (bottom) for chromosome 16 in healthy lymphoblastoids: GM12878 Replicate 1. **c** 250kb bin size Hi-C interaction heatmaps (top) and plots of the first eigenvector from principal component analysis (bottom) for chromosome 16 in ATM mutant fibroblasts: ATM-hTERT Replicate 2 (left) and GM02052 (right).

**Supplementary Figure 8.**
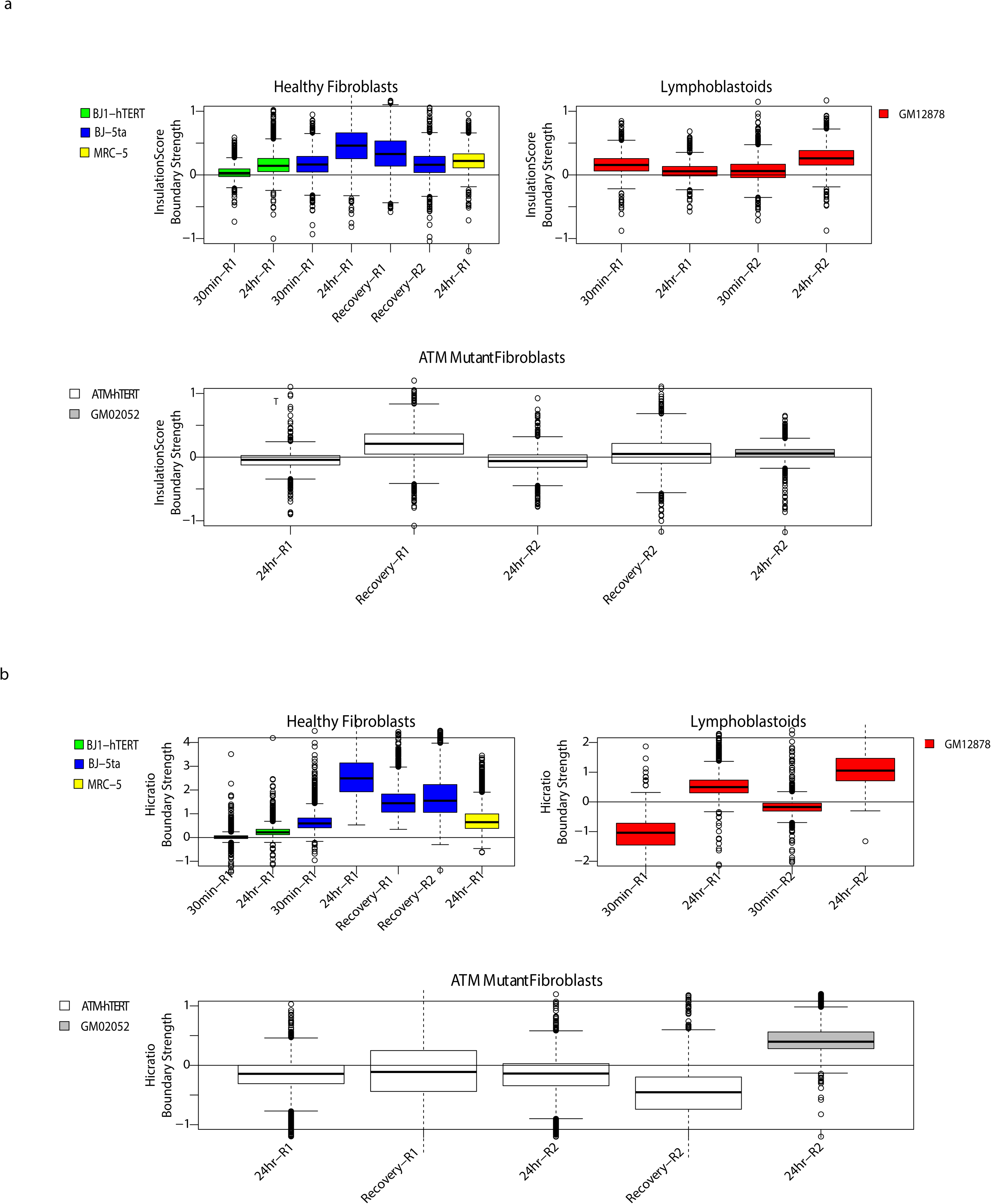
TAD insulation score patterns after exposure to IR are consistent between replicates and among similar cell lines. **a** TAD insulation score boxplots calculated by InsulationScore method for all replicates of all cell lines used in the study. Healthy fibroblast lines: BJ1-hTERT (green), BJ-5ta (blue), and MRC-5 (yellow); Lymphoblastoid line: GM12878 (red); ATM mutant fibroblast lines: ATM-hTERT (white) and GM02052 (gray). **b** Boxplots of TAD insulation scores calculated by Hicratio method for all replicates of all cell lines used in study. Healthy fibroblast lines: BJ1-hTERT (green), BJ-5ta (blue), and MRC-5 (yellow); Lymphoblastoid line: GM12878 (red); ATM mutant fibroblast lines: ATM-hTERT (white) and GM02052 (gray).

**Supplementary Figure 9.**
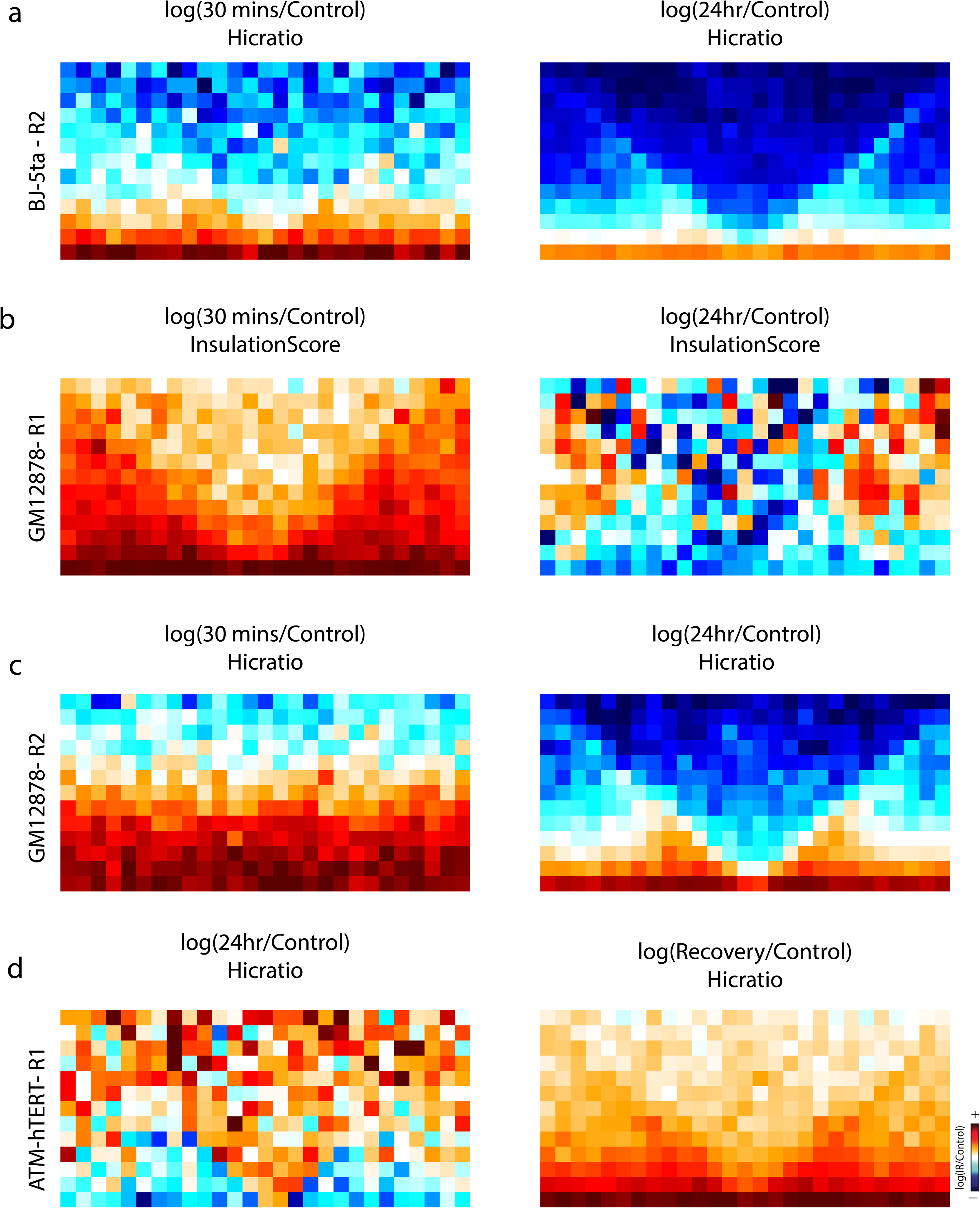
Insulation heatmaps generated using InsulationScore or Hicratio called boundaries. Aggregate contact maps of averaged TAD boundaries called by Hicratio method in **a** BJ-5ta, **c** GM12878, and **d** ATM-hTERT. **b** Aggregate contact map of averaged TAD boundaries called by InsulationScore method in BJ-5ta.

**Supplementary Figure 10.**
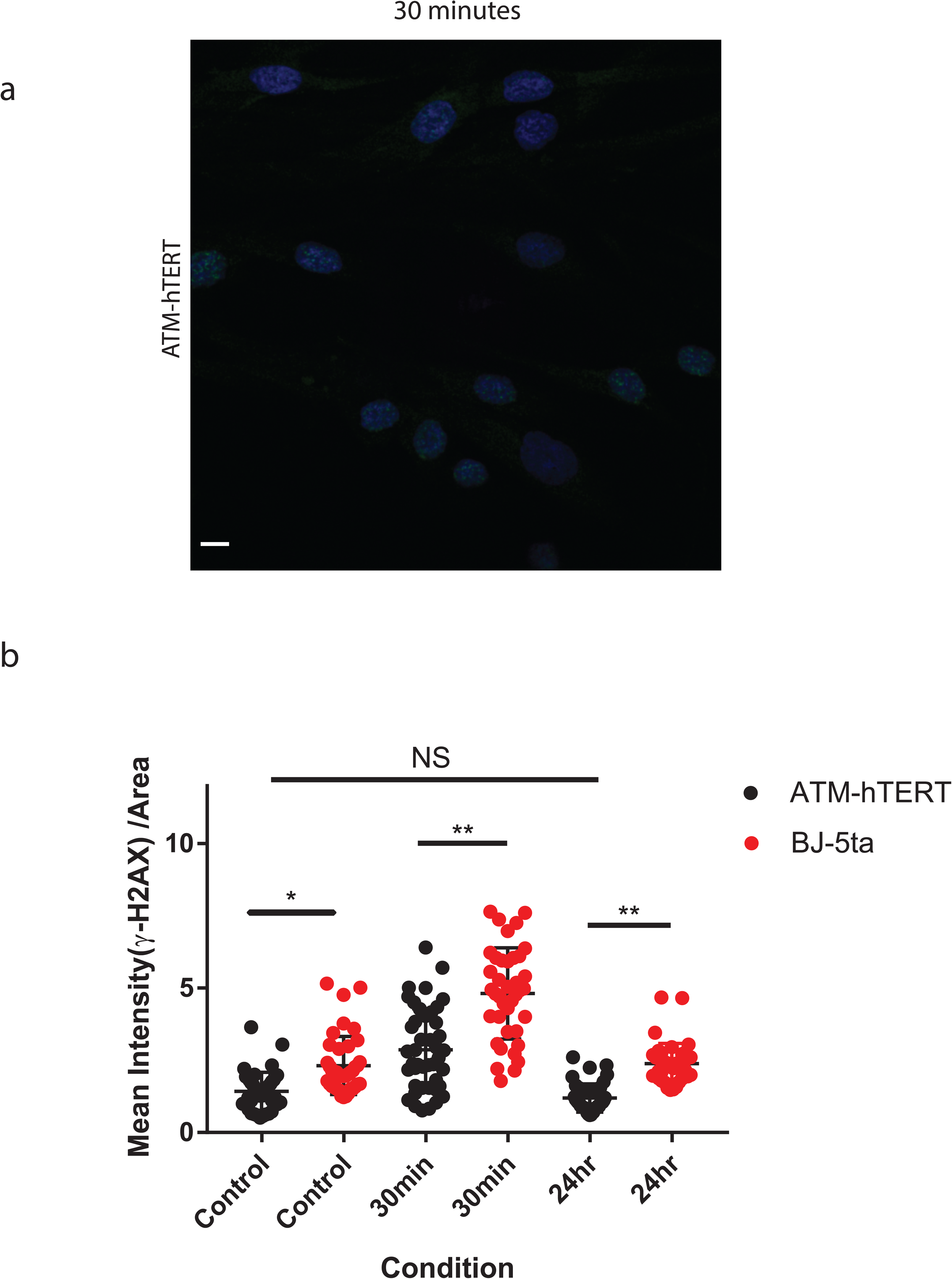
γ-H2AX levels increase in ATM-hTERT cells, but have a weaker response than BJ-5ta. **a** ATM-hTERT fibroblasts stained with yH2AX (green) and DAPI (blue) 30 minutes after exposure to 5 Gy X-rays. Scale bar: 10um. **b** Quantification of mean fluorescence intensity in ATM-hTERT compared to BJ-5ta data in Supplementary Figure 2a (n = 40, * p<0.0020, ** p<0.0001, one way ANOVA). γ-H2AX levels are significantly lower in ATM mutants than BJ-5ta but maintain similar pattern of increase at 30 mins (p <0.0001) and return to initial levels at 24hrs (p > 0.05).

**Supplementary Table 1.**

| Cell Line | Condition | Replicate | Raw # Reads | # Both Sides Mapped | Valid Pairs | Unique Valid Pairs | % Cis | Dangling Ends % |
| --- | --- | --- | --- | --- | --- | --- | --- | --- |
| BJ-5ta | Control | R1 | 180,650,273 | 112,234,560 | 98,769,926 | 62,194,681 | 74.4 | 10.3 |
| BJ-5ta | 30 mins X | R1 | 172,469,514 | 105,572,234 | 101,168,488 | 76,520,452 | 77.9 | 3.8 |
| BJ-5ta | 24 hr X | R1 | 439,993,397 | 294,611,193 | 285,966,444 | 212,551,030 | 86.2 | 1.7 |
| BJ-5ta | Recovery | R1 | 387,242,758 | 247,317,471 | 242,377,386 | 141,227,429 | 86.2 | 1.3 |
| BJ-5ta | Recovery | R2 | 283,878,060 | 173,354,500 | 165,142,153 | 104,333,341 | 77.6 | 4.3 |
| BJ-1 hTERT | Control | R1 | 246,979,150 | 178,487,837 | 166,441,060 | 129,865,596 | 84.4 | 6.2 |
| BJ-1 hTERT | 30 min X | R1 | 231,840,475 | 168,157,410 | 155,628,339 | 125,629,148 | 83.8 | 6.9 |
| BJ-1 hTERT | 24 hr X | R1 | 205,760,134 | 147,470,034 | 134,110,303 | 109,198,540 | 84.8 | 8.4 |
| ATM-hTERT | Control | R1 | 204,830,725 | 168,107,250 | 141,109,197 | 124,465,656 | 78 | 15.8 |
| ATM-hTERT | Recovery | R1 | 389,931,788 | 255,700,038 | 243,088,728 | 125,064,421 | 87.4 | 4.0 |
| ATM-hTERT | 24 hr X | R1 | 179,977,928 | 149,541,854 | 121,284,841 | 111,110,533 | 78.6 | 18.5 |
| ATM-hTERT | Control | R2 | 60,818,985 | 41,342,146 | 34,216,181 | 31,768,043 | 76.9 | 16.9 |
| ATM-hTERT | 24 hr X | R2 | 198,588,290 | 138,204,859 | 110,691,110 | 102,849,684 | 77.8 | 19.5 |
| ATM-hTERT | Recovery | R2 | 98,687,970 | 63,318,090 | 59,208,561 | 48,238,983 | 82.9 | 6.0 |
| GM12878 | Control | R1 | 169,041,926 | 141,427,409 | 111,380,425 | 95,881,352 | 77.7 | 20.9 |
| GM12878 | 30 min X | R1 | 404,673,351 | 254,353,026 | 248,835,472 | 175,309,206 | 80 | 1.4 |
| GM12878 | 24 hr X | R1 | 192,011,866 | 157,657,922 | 122,355,226 | 107,876,878 | 75.2 | 22.0 |
| GM12878 | Control | R2 | 198,416,626 | 141,010,973 | 138,034,618 | 124,040,375 | 77.4 | 1.7 |
| GM12878 | 30 mins X | R2 | 242,425,056 | 156,489,034 | 146,862,286 | 115,111,914 | 79.9 | 5.8 |
| GM12878 | 24 hr X | R2 | 190,188,740 | 136,357,434 | 131,529,834 | 106,197,889 | 83 | 2.6 |
| MRC5 | Control | R1 | 128,868,956 | 105,055,075 | 90,160,337 | 66,947,233 | 73.1 | 14.0 |
| MRC5 | 24 hr X | R1 | 95,372,972 | 77,336,363 | 52,976,419 | 43,945,126 | 82.7 | 31.3 |
| GM02052 | Control | R1 | 402,772,517 | 254,020,052 | 224,921,471 | 154,220,231 | 83.7 | 10.8 |
| GM02052 | 24 hr X | R1 | 384,201,796 | 247,283,552 | 221,226,346 | 139,824,607 | 84.7 | 9.8 |

**Supplementary Table 2.**

| Cell Line | Medium | FBS Content | Additional Supplements |
| --- | --- | --- | --- |
| BJ-5ta | 4:1 DMEM:Medium 199 | 10% | Hygromycin B (0.01 mg/mL) |
| BJ1-hTERT | 4:1 DMEM:Medium 199 | 10% | Hygromycin B (0.01 mg/mL) |
| GM12878 | RPMI 1640 | 15% |  |
| MRC-5 | MEM | 10% |  |
| AG04405 (ATM-hTERT) | DMEM | 15% | 1X Non-essential amino acids |
| GM02052 | MEM | 10% | 1X Non-essential amino acids |

